# Distinct human gut microbial taxonomic signatures uncovered with different sample processing and microbial cell disruption methods for metaproteomic analysis

**DOI:** 10.1101/2020.10.08.331066

**Authors:** Carmen García-Durán, Raquel Martínez López, Inés Zapico, Enrique Pérez, Eduardo Romeu, Javier Arroyo, María Luisa Hernáez, Aida Pitarch, Lucía Monteoliva, Concha Gil

**Author notes:** **Correspondence:** Lucía Monteoliva, Concha Gil.

## Abstract

Metaproteomics is as a promising technique for studying the human gut microbiota, because it can reveal the taxonomic profile and also shed light on the functional role of the microbial community. Nevertheless, methods for extracting proteins from stool samples continue to evolve, in the pursuit of optimal protocols for moistening and dispersing the stool sample and for disrupting microbial cells which are two critical steps for ensuring good protein recovery. Here, we evaluated different stool sample processing and microbial cell disruption methods for metaproteomic analyses of human gut microbiota. An unsupervised principal component analysis showed that different methods produced similar human gut microbial taxonomic profiles. An unsupervised two-way hierarchical clustering analysis identified the microbial taxonomic signatures associated with each method. Proteobacteria and Bacteroidetes identification was favored by moistening the stool samples during processing and by disrupting cells with medium-sized glass beads. Ascomycota identification was enhanced by using large-sized glass beads during sample processing for stool dispersion. Euryarchaeota identification was improved with a combination of small and medium-sized glass beads for cell disruption. Assessments of the relative abundance of Firmicutes, Actinobacteria and Spirochaetes improved when ultrasonication was performed before cell disruption with glass beads. The latter method also increased the overall number of identified proteins. Taxonomic and protein functional analyses of metaproteomic data derived from stool samples from six healthy individuals showed common taxonomic profiles. We also detected certain proteins involved in microbial functions relevant to the host and related mostly to particular taxa, such as B12 biosynthesis and short chain fatty acid production carried out mainly by members in the *Prevotella* genus and the Firmicutes phylum, respectively. Finally, in this metaproteomic study we identified several human proteins, mostly related to the anti-microbial response, which could contribute to determining the beneficial and detrimental relationships between gut microbiota and human cells in particular human diseases.

## 1 Introduction

The human gut microbiota is a complex community of microorganisms that inhabit the human gastrointestinal tract. The gut microbiota comprises nearly 1000 bacterial species, in addition to archaea, fungi, parasites, and viruses (Alarcón et al., 2016; Lloyd-Price et al., 2016). When this ecosystem is balanced and shows high diversity, its close relationship with the host provides beneficial effects, such as the digestion of indigestible foods, defense against pathogens, immunomodulation, and production of vitamins and other beneficial products (Jandhyala et al., 2015). However, even with its great capacity for resilience, the gut microbiota is influenced by many factors, including diet, age, pollution, and the consumption of antibiotics, among others (Jakobsson et al., 2010; David et al., 2014; Jin et al., 2017; Kim and Jazwinski, 2018; Adak and Khan, 2019). These factors can affect its composition and biodiversity, which in some cases, leads to dysbiosis. Although it remains unclear whether dysbiosis is a cause or consequence of host pathological states, the link between pathological states and gut microbiota dysbiosis is a proven fact. Gut microbiota dysbiosis was previously associated with several disease states, including gastrointestinal diseases (Ni et al., 2017; Saffouri et al., 2019), cancer (Wang et al., 2019), Alzheimer’s disease, (Kowalski and Mulak, 2019) and even autism (Fattorusso et al., 2019). Furthermore, the microbiota-gut-brain axis is the link between the gut microbiota and host pathologies related to mental conditions (Wang and Wang, 2016). This link illustrates the importance of the metabolic pathways carried out by gut microbiota, which produce a high diversity of enzymes and metabolites that perform several functions in different parts of the body, not exclusively in the gastrointestinal tract.

Currently, metagenomics is the most common ‘omic’ approach for studying microbiota, due to its ability to provide valuable information about microbial complexity (Jovel et al., 2016). Nevertheless, this strategy lacks the ability to provide functional insights. In this context, metaproteomics is a promising approach for studying gut microbiota, both from the taxonomic point of view and the functional point of view. Indeed, metaproteomics can reveal the main metabolic and functional roles played by the different microorganisms present in gut microbiota (Zhang et al., 2016; Issa Isaac et al., 2019). Moreover, this approach can provide information about proteins that mediate the interactions between the gut microbiota and the host (Blackburn and Martens, 2016). Thus, metaproteomics can provide a better understanding of the functional roles of the gut microbiota, compared to other ‘omics’ approaches (Zhang et al., 2019).

Despite advancements in metaproteomic techniques in recent years (Zhang and Figeys, 2019), the complexity of the microbiota samples, typically derived from stool samples, has made it difficult to devise a suitable protocol for maximizing microorganism recovery. In addition, an optimal microbial cell disruption method is needed to ensure that protein identification is as comprehensive as possible. Because protein extraction methods can affect metaproteomics results (Zhang et al., 2018), the protocol must be optimized to produce representative taxonomic profiles and identify the most significant metabolic roles carried out by gut microbiota.

In addition to optimizing the extraction of microbial proteins from a stool sample, bioinformatics is a crucial step in the metaproteomics workflow. The use of specific software, such as Metalab (Cheng et al., 2017), has facilitated the analysis of the large amounts of data generated in metaproteomics approaches for studying gut microbiota. This software accesses a protein dataset that was created from more than 1000 human microbiota samples, which allows a high number of peptide/protein identifications. The software also allows functional characteristics to be assigned to the identified proteins at specific taxon levels.

In this study, we aimed to optimize the isolation and characterization of human gut microbiota by analyzing one stool sample with different protocols that combined different sample processing and microbial cell disruption methods. We assessed the relevance of the optimized technique by characterizing and comparing six human gut microbiome samples, according to their taxon profiles and the functions of the identified proteins in each individual sample.

## 2 Materials and methods

### 2.1 Stool samples

Stool samples were collected from 6 healthy adult volunteers with their informed consent. The 6 samples belonged to 3 females and 3 males, aged between 31 and 52 and were named S1 to S6. None of them had been under antibiotics treatment during the previous year to sampling. Only one subject reported gastrointestinal problems within 3 months prior to sampling. Feces were stored at −80°C until processing.

### 2.2 Metaproteomic analysis of stool samples

We combined two different stool sample processing (SSP) and four cell disruption methods (CDM). All these protocols are schematized in Figure 1.

**Figure 1.**
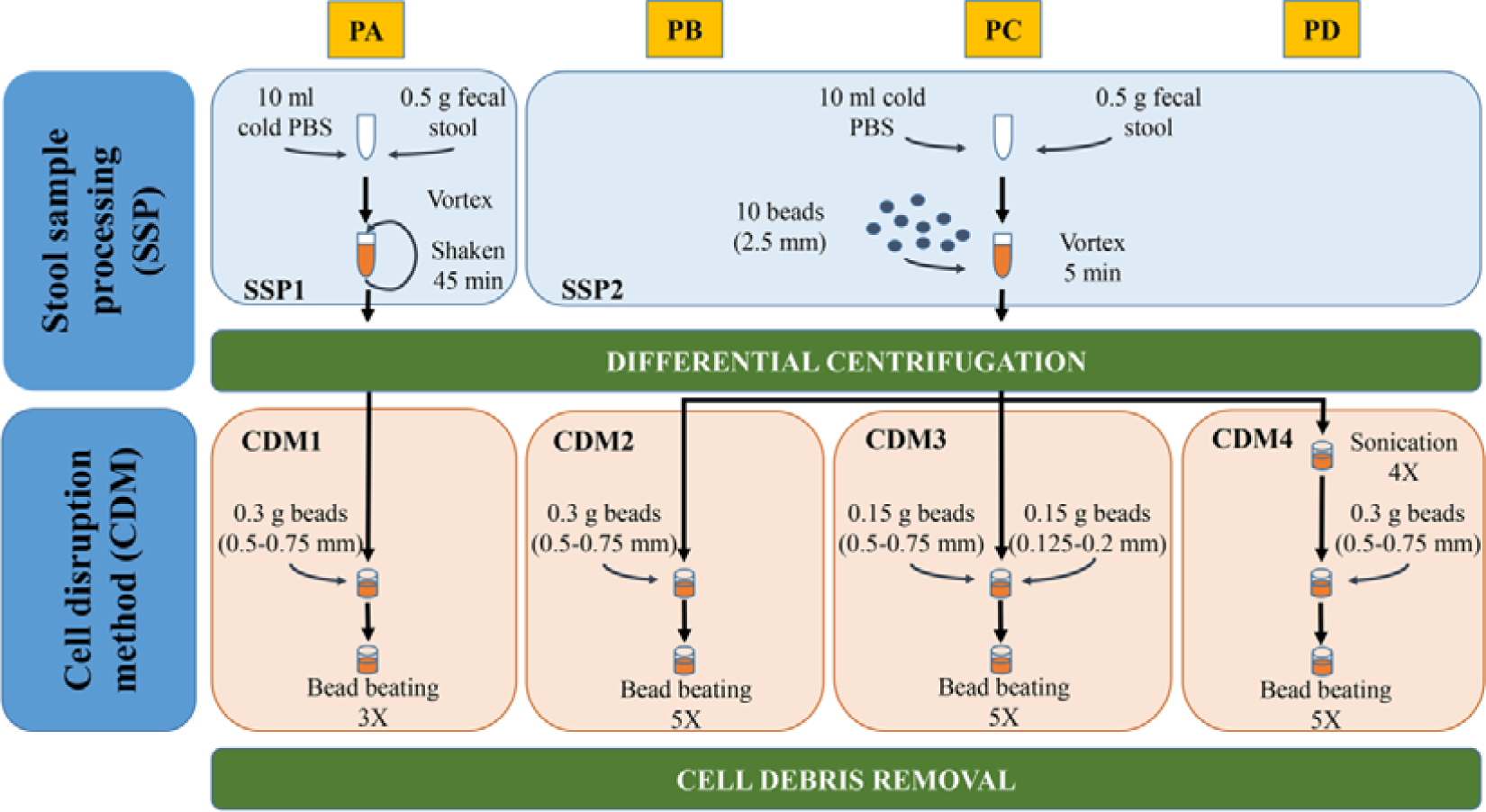
Schematic representation of the different SSP and CDM combinations carried out with stool sample S1 (PA-PD). For more details see materials and methods and results sections.

#### 2.2.1 Microbial enrichment by SSP

SSP1 was modified from different works (Tanca et al., 2015; Zhang et al., 2018). Briefly, 0.5 g of the stool sample were resuspended in 10 mL ice-cold phosphate buffer (PBS), vortexed and shaken in a tube rotator for 45 min at 4 °C. The hydrated samples were subsequently subjected to low speed centrifugation at 500 g, 4 °C for 5 min which favored the sedimentation of particulate and insoluble material allowing the collection of the supernatant (containing microbial cells). The supernatant was then transferred to a new tube and kept at 4 °C while the pellets were resuspended in 10 mL ice-cold PBS, repeating this process two other times. Finally, the supernatants (∼30 mL) were centrifuged at 14 000 g and 4 °C for 20 min to pellet the microbial cells. Pellet was then washed three times with PBS. The washing step was performed by resuspension of cell microbial pellet in ice-cold PBS followed by centrifugation at 14 000 g and 4 °C for 20 min. The resulting washed pellet was subjected to protein extraction. SSP2 is a variation of SSP1 in which the 45 min shaking step in the tube rotator was changed for a 5 min vortexing step with 10 glass beads (2.5 mm).

#### 2.2.1 CDM for protein extraction

Four different CDM (1-4) were carried out for cell disruption and protein extraction. CDM1 was modified from Zhang et al. (2018). Briefly, the microbial pellet was resuspended in 500 μL lysis buffer (4% SDS (w/v) in 50 mM Tris-HCl buffer pH 8.0) followed by incubation in a Thermomixer Comfort (Eppendorf) at 95 °C for 10 min with agitation. After cooling, the lysates were transferred to a 2-mL screw-cap tube containing 0.3 g glass beads (0.5-0.75 mm) (Retsch). Bead beating was carried out using a FastPrep-24 machine (MP Biomedicals Inc., USA) at a speed of 6.5 ms^−1^ for 90 s (3 rounds 30 s each with 5 min interval on ice). Finally, cell debris and beads were removed through centrifugation at 16 000 g and 4°C for 10 min. The supernatant was transferred into a new tube and centrifuged again at 16 000 g and 4°C for 10 min to remove any remaining particulate debris. The final supernatant was used for protein quantification and purification. CDM2 was performed in the same way to CDM1 but increasing to 5 the rounds of bead beating. CDM3 was performed similar to CDM2 but introducing two different sizes of glass beads (0.15 g of 0.5-0.75 mm and 0.15 g of 0.125-0.2 mm). CDM4 was modified from Zhang et al. (2017). Briefly, the microbial pellet was resuspended in 500 μL 4% SDS (w/v) lysis buffer followed by incubation in a Thermomixer Comfort at 95 °C for 10 min with agitation. After cooling and once the lysates were transferred to a 2-mL screwcap tube, 4 ultrasonications (30 s each with 1 min interval on ice) using Vibra Cell Sonicator (Sonics & Materials Inc., USA) with an amplitude of 40% were carried out prior to bead beating. After sonication, bead beating conditions (including beads sizes, speed and rounds) and supernatant collection of soluble proteins were performed as described in CDM2. Combination of the different SSP and CDM resulted in four protocols named A-D (Figure 1).

### 2.3 Peptide sample preparation for mass spectrometry

Proteins were precipitated using methanol/chloroform to remove SDS, and then resuspended in 8 M urea for in-solution trypsin digestion. The samples were quantified with Qubit 3.0 (ThermoFisher Scientific) and 50 µg proteins were reduced with 10 mM dithiothreitol (DTT) at 37 °C for 45 min and alkylated with 55 mM iodoacetamide (IAA) at room temperature for 30 min in darkness. 1/25 trypsin (w/w) (Roche Molecular Biochemicals) was added to each sample in 25 mM ammonium biocarbonate for trypsin digestion at 37 °C overnight. The peptides from proteins digested were desalted and concentrated with C18 reverse phase chromatography (OMIX C18, Agilent technologies) and peptides were eluted with 80% acetonitrile (ACN) / 0.1% trifluoroacetic acid. The eluent was then freeze-dried in Speed-vac (Savant) and resuspended in 12 µL 2% ACN / 0.1% formic acid (FA). Finally, peptides were quantified in Qubit fluorimeter (Thermo Scientific) and 1 µg peptides were used for LC-MS/MS analysis.

### 2.4 Liquid chromatography-tandem mass spectrometry analysis

One µg peptides were loaded for RP-nano-LC-ESI-MS/MS analysis on an EASY-nLC 1000 System (Proxeon) coupled to a Q Exactive HF mass spectrometer (Thermo Scientific Inc.). The peptides were ionized by electrospray in positive mode in DDA (data-dependent acquisition). From each MS scan (between 350 to 2 000 Da) the 15 most intense precursors (charge between 2+ and 5+) were selected for their HCD (high collision energy dissociation) fragmentation and the corresponding MS/MS spectra were acquired. Peptides were eluted using 240 min gradient.

The mass spectrometry proteomics raw data have been deposited to the ProteomeXchange Consortium (http://www.proteomexchange.org) via PRIDE partner repository with the database identifiers PXD020786 and PXD020787.

### 2.5 Bioinformatics analysis

For data processing, MetaLab version 2.0 was used (Cheng et al., 2017). This software provided peptide and protein identification and quantification, taxonomic profiling and functional annotation. Briefly, the human gut microbial gene catalog with 9 878 647 sequences (available on http://meta.genomics.cn/) was employed for generating a reduced, sample-specific protein database (Li et al., 2014). The resulting database was used for the peptide characterization using the Andromeda search engine from MaxQuant version 1.6.10.43 which is integrated in MetaLab. Quantitative analysis was performed employing the maxLFQ algorithm in MaxQuant. Proteins identified by the same set or a subset of peptides were grouped together as one protein group. Human peptides were identified using Uniprot DB (www.uniprot.org) (Consortium, 2019) restricted to human taxonomy (downloaded 02.18.2020 with 74 451 sequences) as described previously for microbial peptides characterization. Taxonomy analysis of the generated peptide list was performed matching peptides to the lowest common ancestor (LCA). Abundance data of both peptides and taxa were provided according to the LFQ intensities. The quantitative information of each taxon, which was calculated as the sum intensity of all distinctive peptides of that taxon, was used to estimate its relative abundance. The taxonomic results were manually filtered to not retain human taxa and only taxa identified with at least 3 peptides were considered.

Functional annotation for the identified proteins was also assigned and automated generated from MetaLab to directly obtain information about COG and KEGG. For an in deep functional analysis, KO (KEGG Orthology) database information (molecular functions as functional orthologs defined in the context of KEGG molecular networks) was used (http://meta.genomics.cn). Taxon-function analysis was carried out using taxonomic information of the enrichment analysis from iMetaLab platform (http://shiny.imetalab.ca/) in combination with KO database information.

KEGG pathway was downloaded from KEGG website (http://www.kegg.jp). STRING analysis was done with STRING program (version 11.0). Graphs were made with Infogram (https://infogram.com/).

### 2.6 Statistical analysis

Unsupervised principal component analysis (PCA) was carried out to group the stool samples from one selected healthy individual (S1) extracted with the different established protein extraction protocols on the basis of the similarities in their gut microbial taxonomic patterns observed from the metaproteomic data. The degree of homology or relative similarity of these profiles among the study groups was assessed with the t-test, whereas their degree of homogeneity or relative variation within each study group was evaluated by analysis of variance (ANOVA). Unsupervised two-way hierarchical clustering analysis (HCA) was performed to cluster the stool samples extracted with the four protocols and different identified (phylum, family or genus) taxa simultaneously according to the similarities in the gut microbial taxonomic profiles of each extraction method and the pattern of each identified gut microbial taxon across all protein extraction protocols, respectively. Relative abundance levels were normalized by mean centering the gut microbial taxonomic profiles for each protein extraction protocol and then by mean centering each pattern of each identified microbial taxon across the four protocols. Statistical significance was set at p < 0.05 (two-sided).

### 3 Results and discussion

#### 3.1 Protein extraction protocol effects on protein yield and identification

In metaproteomics, protein identification yield depends on the methods used for protein isolation and solubilization, particularly in complex samples, such as stool samples (Zhang et al., 2018). We used one out of six stool samples collected (S1) to examine the effects of four different protein extraction protocols (referred to as PA, PB, PC, and PD) on the metaproteomic analysis of gut microbiota (Figure 1). These protocols included two main steps: (*i*) stool sample processing (SSP), which is needed to disperse the feces to harvest the microbial cells, and (*ii*) a cell disruption method (CDM) for breaking up microbial cells to isolate and solubilize their proteins. In PA, the SSP was performed with three rounds of 45-min shaking/low speed centrifugation (SSP1). In PB, PC, and PD, a faster, easier SSP was devised by adding glass beads and vortexing for 5 min (SSP2). We also compared different CDMs by testing different sizes of glass beads (PC) and an additional sonication step (PD).

When we compared PA and PB, which differenced by the SSP (Figure 1), we observed that PA provided a higher number of peptide-spectrum matches (PSMs) and more peptide and protein identifications than PB (Table 1). This result indicated that the longer treatment was important. With three 45-min shaking rounds at 4°C, PA moistened the stool sample correctly, which allowed the recovery of as many microbial cells as possible. Nevertheless, we obtained even higher numbers of peptide sequences and protein groups with PD. This result indicated that the ultrasonication in the CDM step contributed to increasing protein recovery from the gut microbiota. Furthermore, PC showed higher numbers of peptide and protein identifications than PB. Therefore, the use of different sizes of glass beads seemed to facilitate the disruption of microbial cells, but not to the extent observed with ultrasonication.

**Table 1.**
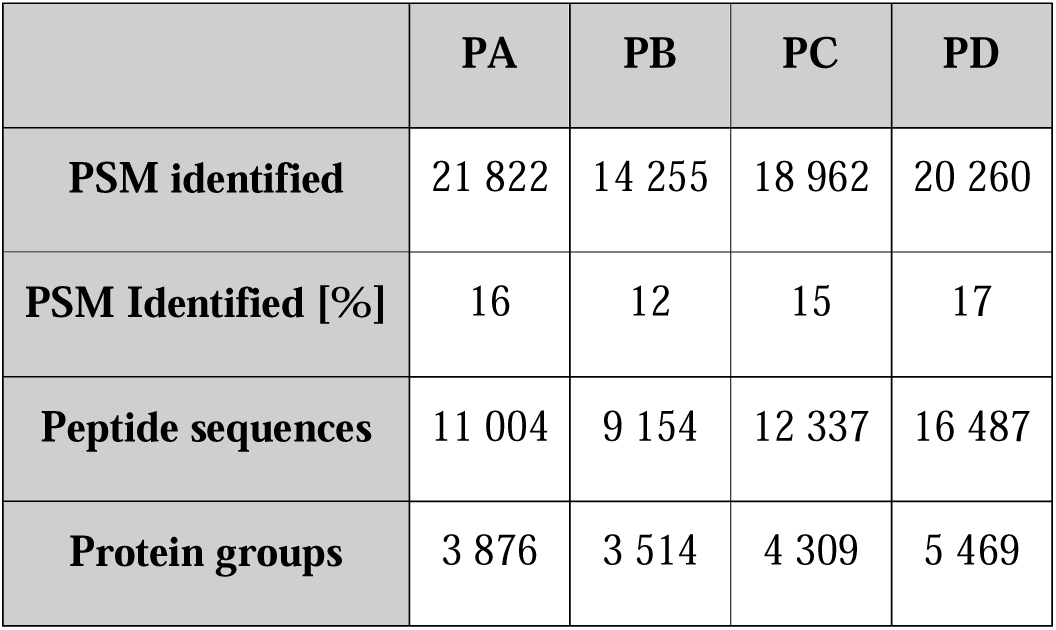
Metrics of metaproteome analysis of S1 using protocols A-D.

|  | PA | PB | PC | PD |
| --- | --- | --- | --- | --- |
| **PSM identified** | 21 822 | 14 255 | 18 962 | 20 260 |
| **PSM Identified [%]** | 16 | 12 | 15 | 17 |
| **Peptide sequences** | 11 004 | 9 154 | 12 337 | 16 487 |
| **Protein groups** | 3 876 | 3 514 | 4 309 | 5 469 |

#### 3.2 Impact of protein extraction methods on taxonomic profiles of human gut microbiota

Next, we explored the influence of the four protein extraction protocols on the ability to identify the taxonomic composition of the gut metaproteome in S1. An unsupervised PCA showed that the taxonomic profiles at the phylum, family, and genus levels were similar among the four gut microbial protein extraction methods (Figure 2A). However, the second principal component revealed that 0.87-1.10% of the dataset variance could be explained by differences between SSP1 and SSP2. Consistent with the PCA results, an unsupervised two-way HCA uncovered two major clusters that identified separate taxonomic profiles of the gut metaproteome in S1. The taxonomic profiles obtained with SSP1 (PA) formed a single cluster, and the profiles obtained with SSP2 (PB, PC, and PD) formed a different discrete cluster at the phylum, family, and genus levels (Figure 2B).

**Figure 2.**
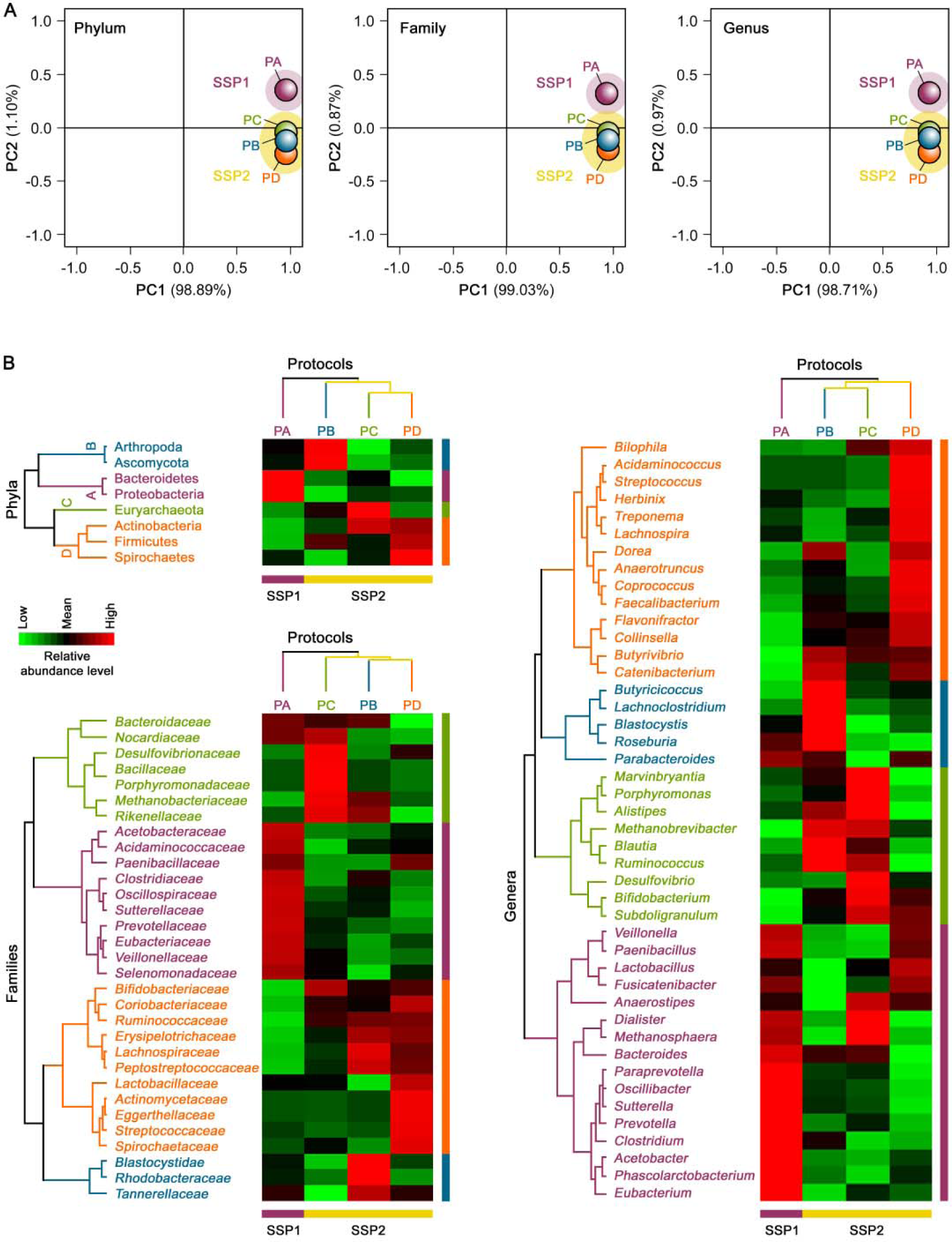
Unsupervised (A) PCA and (B) HCA of the taxonomic profiles of the gut metaproteome of S1 isolated with different microbial protein extraction protocols.

Each protein extraction protocol (PA, PB, PC, and PD) was associated with a specific gut microbial taxonomic signature (subclusters A, B, C, and D, respectively, Figure 2B) at the phylum level. In turn, each taxonomic signature was related to one of the three life domains or superkingdoms (Bacteria, Archaea, and Eukarya). The PA-associated signature (subcluster A) comprised the bacterial phyla, Proteobacteria and Bacteroidetes. This result supported the notion that SSP1 and medium-sized glass beads for cell disruption could favor protein extractions from Gram-negative bacteria in the human gut microbial community. The PD-associated signature (subcluster D) included the bacterial phyla, Firmicutes, Actinobacteria, and Spirochaetes. This finding highlighted the effect of an additional ultrasonication step before mechanical cell disruption with medium-sized glass beads. Thus PD facilitated gut microbial protein extractions from Gram-positive bacteria, which are more difficult to lyse than Gram-negative bacteria (Dridi et al., 2009; Salonen et al., 2010; Zhang et al., 2018). In addition, PD enhanced protein extractions from spirochetes, which have a unique cell envelope due to their endoflagella. These Gram-negative bacteria have been shown to share some Gram-positive bacteria characteristics, unlike other Gram-negative bacteria. The PC-associated signature (subcluster C) displayed a significantly higher proportion of the archaeal phylum, Euryarchaeota. PC also improved gut microbial protein extractions from the bacterial phylum, Actinobacteria. These findings revealed that the combination of small and medium-sized glass beads added during intensive mechanical cell disruption could enhance protein extractions of hard-to-lyse gut microbiota members, such as methanogenic archaea and some Gram-positive bacteria, consistent with findings in previous studies (Dridi et al., 2009; Salonen et al., 2010; Zhang et al., 2018). In contrast, the PB-associated signature (subcluster B) integrated the eukaryal phyla, Ascomycota and Arthropoda. These data indicated that protein extractions from eukaryotic microorganisms in the human gut microbiota could be improved with SSP2 and medium-sized glass beads, consistent with previous studies (Pitarch et al., 2008; Pitarch et al., 2009). Remarkably, PB also recovered a high proportion of the dominant gut microbial phylum, Firmicutes, but the proportion was lower than the proportion recovered with PD.

Our results also suggested that moistening the stool samples during SSP (SSP1) could be the key factor for studying Proteobacteria in the human gut microbial community. The members of this phylum are present in low abundance in the natural human gut microbiota, but they can serve as a potential diagnostic microbial signature of gut microbiota dysbiosis and disease risk (Shin et al., 2015). Therefore, PA could be the method of choice for metaproteomics studies related to the human gut microbiota in health and disease states.

#### 3.3 Metaproteomics analysis of gut microbiota from healthy adults

We carried out a metaproteomic analysis of gut microbiota in stool samples from 6 healthy adults (S1-S6) with PA. Upon characterizing microbial biodiversity, we focused on the contribution of each taxon to the metabolic processes and cell functions involved in the human gut environment, which could be linked to specific states of the host. We identified 154 246 PSMs, 37 080 peptide sequences, and 10 686 protein groups. Among the 6 samples, the means were 25 503 PSMs, 11 712 peptide sequences, and 4253 protein groups (Table 2). Unlike taxonomic genomic studies, which are based on 16S rRNA, metaproteomic analyses allow the identification of human proteins. Nevertheless, for taxonomic purposes, we filtered out the human proteins and focused exclusively on non-human peptides. Considering only proteins with at least 3 distinctive peptides, the identified proteins corresponded to 11 phyla, 19 classes, 25 orders, 34 families, 53 genera, and 105 species (Table S1). We could not assign 17% of the identified peptides to any phylum. However, among all identified peptides, 27% could be assigned to species level. The percentage of peptides that could be assigned to superior taxon levels increased as the level increased: we assigned 54% at the genus level, 56 % at the family level, 71% at the order level, and 72% at the class level.

**Table 2.** Metrics of metaproteome analysis of healthy adult stool samples (S1-S6).

|  | S1 | S2 | S3 | S4 | S5 | S6 | Mean |
| --- | --- | --- | --- | --- | --- | --- | --- |
| PSM identified | 20 915 | 28 240 | 25 146 | 29 509 | 22 374 | 28 062 | 25 503 |
|  | 154 246* |  |  |  |  |  |  |
| PSM identified [%] | 15 | 20 | 18 | 20 | 16 | 20 | 18 |
| Peptide sequences | 10 198 | 12 866 | 10 975 | 13 222 | 10 544 | 12 855 | 11 712 |
|  | 37 080* |  |  |  |  |  |  |
| Protein groups | 3 926 | 4 602 | 3 732 | 4 525 | 4 110 | 4 722 | 4 253 |
|  | 10 686* |  |  |  |  |  |  |

\*Total number of PSM, peptides sequences and protein groups identified in the study.

#### 3.3.1 Taxonomic analysis of gut microbiota

Bacteria was the most abundant superkingdom in the human gut microbiome; we found that it represented 96-99% of the microbiome, consistent with previous studies (Kolmeder et al., 2016; Tanca et al., 2017). In agreement with other works (Arumugam et al., 2011; Kolmeder et al., 2012; Kolmeder et al., 2016; Tanca et al., 2017; Zhang et al., 2017; Zhong et al., 2019), the four most abundant phyla in our 6 samples were Firmicutes, Bacteroidetes, Proteobacteria, and Actinobacteria (Figure 3A). The dominant phylum was Firmicutes, except in S3, where Bacteroidetes was the principal phylum. Proteobacteria is an important phylum in diseases states. It is normally found in very low abundance in healthy individuals. Accordingly, for studies on effects of the microbiome in diseases, it could be key to choose a convenient protocol that allows adequate recovery of Proteobacteria proteins to assess and compare their abundance in normal and disease states.

**Figure 3.**
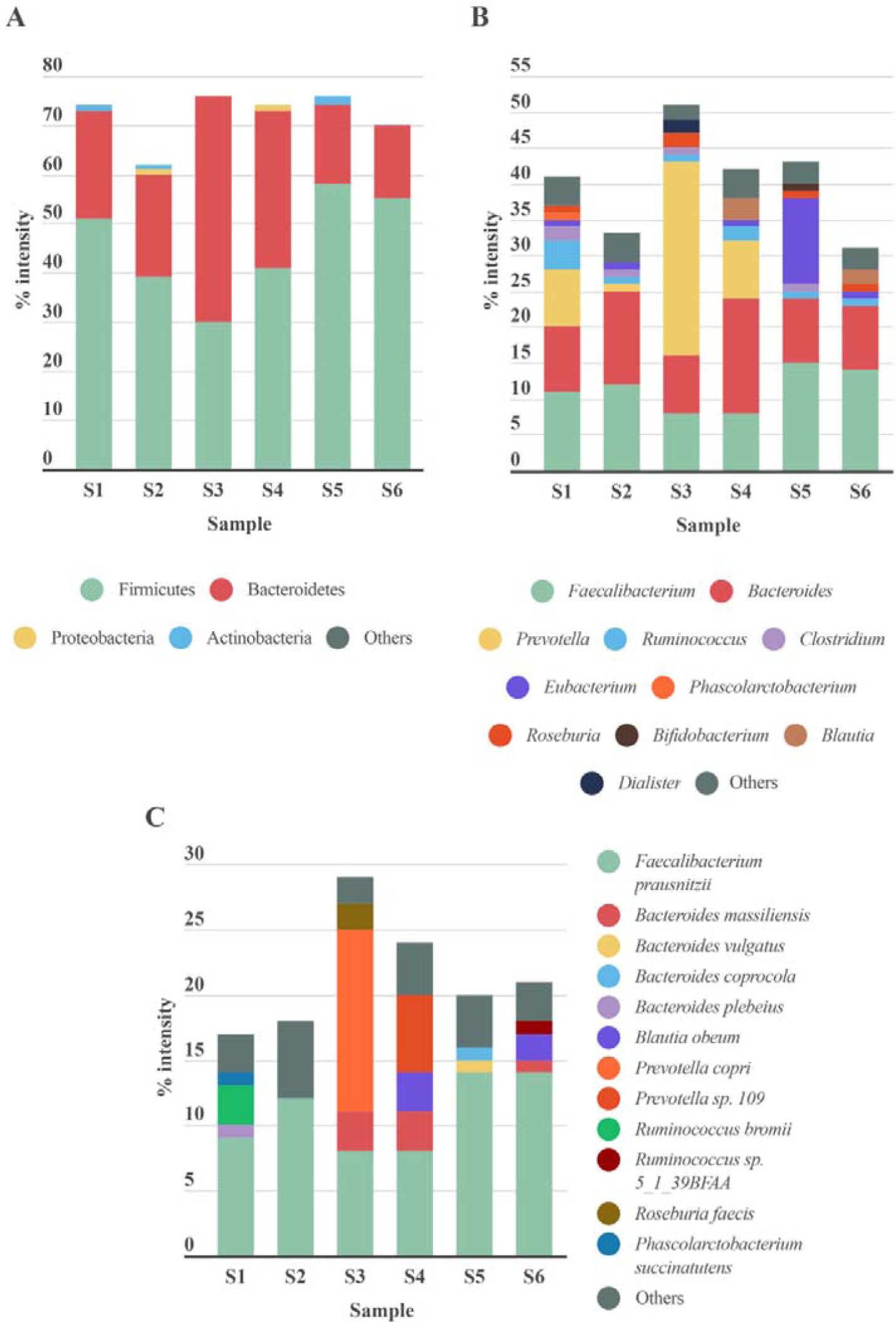
Taxonomic distribution of gut microbiota detected in stool samples from 6 healthy adults (S1-S6). (A) Phylum, (B) genus l and (C) species levels. Abundance of each phylum/genus/species is correlated to the percentage of peptide intensity ([sum intensity of peptides of the phylum]/[total intensity of sample]). Only those taxon categories with >1% of abundance are represented in the graphs. The rest are included in the‘other’ category.

The abundance of the Archaea superkingdom was less than 0.5% in our samples. Euyarchaeota was the only phylum we identified in this superkingdom (Table S1). In the fungi kingdom, we found the Ascomycota phylum, but it was only present in 3 out of the 6 samples, and its mean abundance was 0.01% (Table S1).

The most abundant genera were *Faecalibacterium* (phylum Firmicutes) and *Bacteroides* (phylum Bacteroidetes) (Figure 3B). *Ruminococcus*, another genus from Firmicutes, was highly abundant in S1 and S4, and similarly, *Blautia* was abundant in S4 and S6. S1, S4, and particularly S3, were enriched in the genus *Prevotella* (phylum Bacteroidetes), which represented more than 8% in S1 and S4, and up to 27% in S3. A reciprocal pattern was reported for the abundances of *Prevotella* and *Bacteroides* (both from phylum Bacteroidetes) (Hoffmann et al., 2013), consistent with our results. The high abundance of *Prevotella* has been associated with a high consumption of fiber (David et al., 2014), and *Bacteroides* was linked to the Western diet, based on the relatively high meat consumption (Wu et al., 2011). Therefore, these differences in abundance among the samples could be partly explained by differences in diet among the corresponding individuals. In S3, we found *Akkermansia* present at an abundance of 0.04 % (Table S1), consistent with a previous study (Kolmeder et al., 2012). A species of *Akkermansia* genus, *A. muciniphila*, is a mucin degrading bacteria. This species is gaining interest, because it induces several host responses, due to its proximity to the mucus layer. Indeed, *A muciniphila* was associated with glucose control, inflammation, and improving the gut barrier (Everard et al., 2013). S5 showed an elevated abundance of *Eubacterium:* the abundance was 12% in S5, but less than 2% in the other samples (Figure 3B). Consequently, it would be interesting to study the function of gut *Eubacterium* in more detail. In the Actinobacteria phylum, *Bifidobacterium* was the main genus in our 6 samples; the average abundance was below 4 %, consistent with previous studies (Riviere et al., 2016). Depending on the sample studied, we observed different compositions of taxa in the Proteobacteria phylum. This phylum is normally associated with diseases states and gut microbiota dysbiosis (Rizzatti et al., 2017). The role of this phylum in abnormal physical states has been widely studied, but its contribution to normal conditions is less well known. The main genus in most of the samples was *Sutterella*, but the main genus in S4 and S6, was *Bilophia. Sutterella* spp. have been related to several gastrointestinal disorders, but it is thought that they only have a mild-proinflammatory capacity. However, they keep the immune system alert, due to their ability to adhere to intestinal epithelial cells (Hiippala et al., 2016). In this genus, the *B. wadsworthia* was the only species which was identified in our study. This bacterium was associated with fat-rich diets and intestinal inflammation. Emerging studies have suggested that limiting the prevalence of *B. wadsworthia* in the gut microbiome could have a therapeutic effect on metabolic diseases and intestinal inflammation (Natividad et al., 2018). S2 was the only sample enriched in *Desulfovibrio*. In S1, the taxa composition was more variable, with high abundances of *Acetobacter* and *Azospirillum*, in addition to *Sutterella*.

The Euryarchaeota phylum was only represented by the *Methanobrevibacter* genus (0.06%), and it was only present in S1. However, this finding highlighted the relatively high abundance of this genus within the Archaea gut microbiome (Hoffmann et al., 2013; Lloyd-Price et al., 2016).

At the species level (Figure 3C), consistent with previous studies (Kolmeder et al., 2012; Zhang et al., 2017; Lin et al., 2018), we found that the main species was *Faecalibacterium prausniizii* (8-14%), except in S3, where *Prevotella copri* had an abundance of 14%. Previous studies showed that *F. prausnitzii* displayed beneficial effects against different alterations in the gastrointestinal tract (Bjørkhaug et al., 2019), and it had anti-inflammatory properties (Debyser et al., 2016). In the other hand, the role of *P. copri* in human health remains unclear. The presence of this species in the gut (more common in non-westernized populations) varies largely between individuals (Tett et al., 2019), but when it is present, it is normally the most abundant species (Arumugam et al., 2011), consistent with our findings. *P. copri* was associated with inflammation and a higher risk of developing rheumatoid arthritis (Scher et al., 2013), even though its presence in the gut microbiome was also linked to a healthier diet. To explain these different behaviors, it might be very important to analyze the strain-level diversity and how it is influenced by the diet. Different strains were found in Western and non-Western populations, and these strains had different functions (De Filippis et al., 2019). The occurrence of this species was related to a reduction in the abundance of *Bacteroides* spp. (Franke and Deppenmeier, 2018), however this was not observed in our work. The proportion of *Bacteroides spp* in S3 was not smaller than the proportions observed in the other samples (Figure 3C), despite the fact that the *Bacteroides* genus comprised a smaller proportion of the genera in S3, compared to the other samples (Figure 3B). Regarding *Bacteroides*, different species and abundances were observed among the 6 samples. *B. plebeius* was the major representative in S1, and *B. massiliensis* was the major representative in S3, S4, and S6. In S5, *B. vulgatus* and *B. coprocola* were the main representatives of the *Bacteroides* genus.

#### 3.3.2 Functional characterization of microbial proteins

Focusing on a functional analysis, we used Metalab to obtain information about the different functional assignments of the identified proteins (Cheng et al., 2017). In this metaproteomic approach, 89.5%, 64.7% and 61.5% of the proteins identified could be assigned to entries from the COG, KEGG, and KO databases, respectively. We first calculated the total protein intensity that corresponded to a specific function, by summing the intensities of all proteins associated with the function. Then, we estimated the percentage of each function, when the total intensities of all identified proteins were set to 100%.

An enrichment analysis showed that the main core COG category functions in all 6 samples were: ‘translation, ribosomal structure and biogenesis’, ‘carbohydrate transport and metabolism’, and ‘energy production and conversion’ consistent with previous studies (Debyser et al., 2016). These functions represented nearly 40% of the total protein intensity (Figure 4). This microbiota carbohydrate metabolism could facilitate the host exploitation of different food sources that would otherwise be indigestible. For example, S1 showed a high abundance of *Ruminococcus:* nearly 98% of all glycosidases identified in this sample belonged to members of this genus. This finding was consistent with the great capacity of *Ruminococcus* in carbohydrate degradation described previously (La Reau et al., 2016). We also found that other functions were highly represented in only some samples; for example, ‘cell motility’ was highly represented in S5, and ‘coenzyme transport and metabolism’ was highly represented in S3 (Table S2).

**Figure 4.**
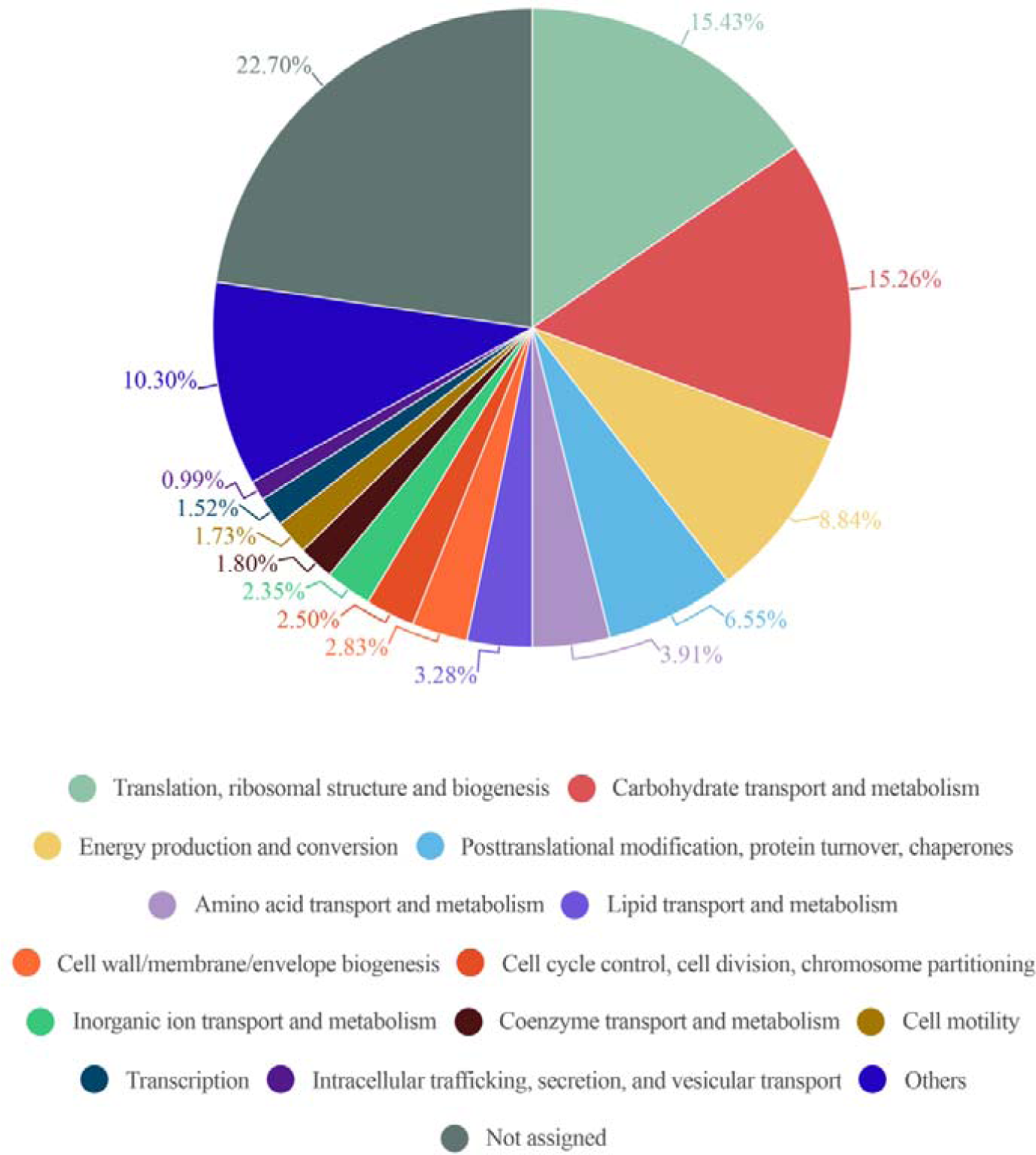
Circle plot representation of the abundance of COG functional entries displayed by the identified gut microbiota proteins from 6 healthy adults. Abundance in each sample is calculated as the percentage resulting from summing up all intensities belonging to proteins annotated to a specific COG category in this sample/total intensity of all proteins annotated to any COG category in this sample. The mean of the COG category abundance of the 6 samples is represented.

The most abundant protein observed in our 6 samples was the glycolytic protein, glyceraldehyde-3-phosphate dehydrogenase (Table S3). In addition, glutamate dehydrogenase, which was the most abundant intestinal protein in another study (Kolmeder et al., 2012), was in the top 30 proteins found in ours samples.

From data obtained from Metalab software we can match taxonomy with the functionality of each identified protein. Therefore, we could assign specific functions to specific taxa. As expected, a general analysis based on COG information, showed that proteins involved in the main functions belonged to the more abundant phyla. However, some specific functions were linked to a specific phylum (Figure 5). For example, ‘lipid transport and metabolism’ was, mostly represented by Firmicutes proteins. In contrast, functions like ‘inorganic ion transport and metabolism’, ‘coenzyme transport and metabolism’, ‘intracellular trafficking, secretion, and vesicular transport’, ‘extracellular structures’ and ‘cell cycle control, cell division, and chromosome partitioning’ were mostly linked to Bacteroidetes. The high number of functions associated with Bacteroidetes could be due to its taxonomic diversity.

**Figure 5.**
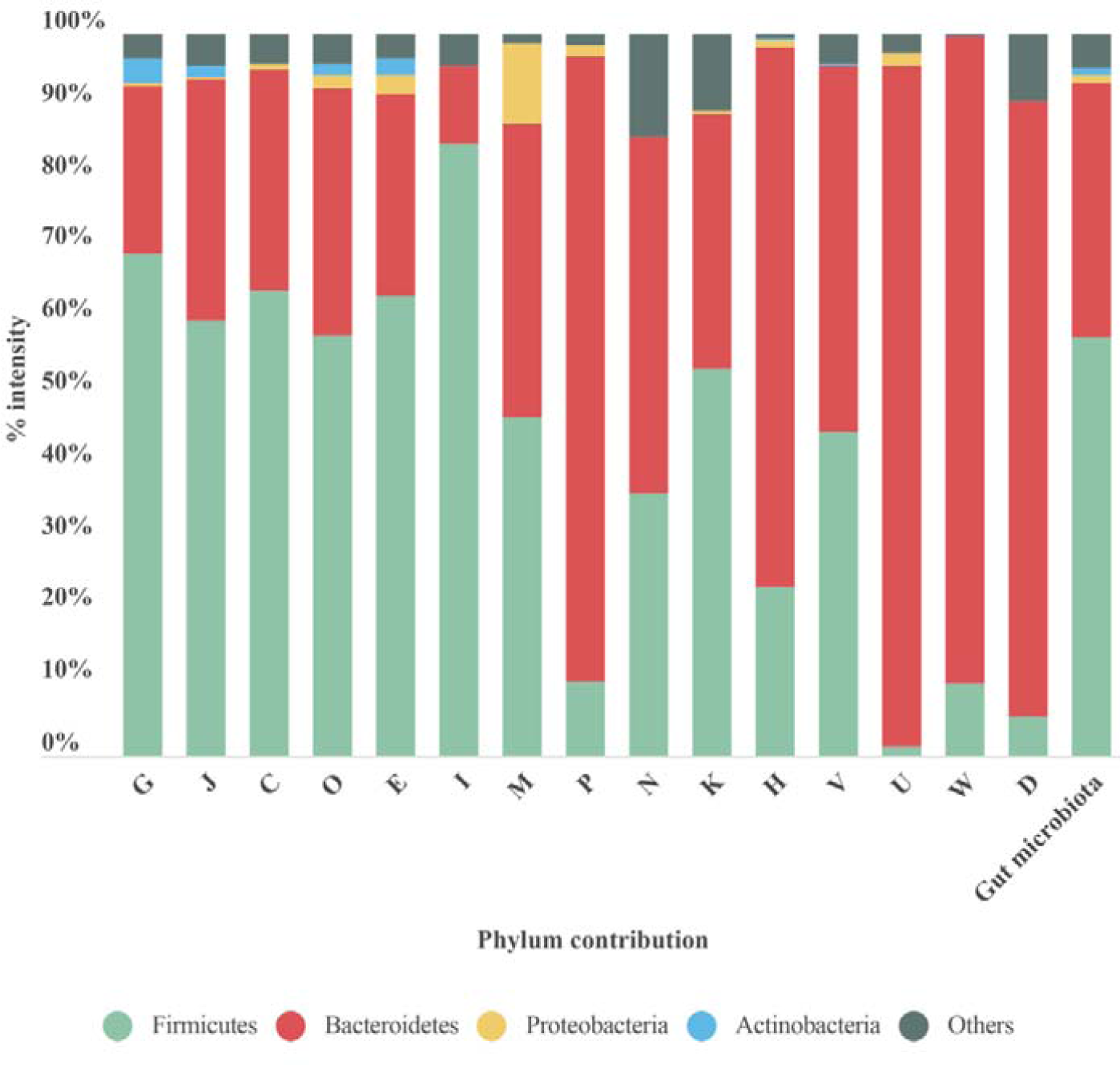
Bar plot representing the contribution of each phylum to the COG category analyzed. The last bar represents the taxonomic distribution of these phyla in the gut microbiota samples. Each phylum’s conribution in each sample is calculated as the sum intensity of proteins of this phylum annotated to the COG category. This bar plot represents the mean abundance of the 6 samples. G, Carbohydrate transport and metabolism; J, Translation, ribosomal structure and biogenesis; C, Energy production and conversion; O, Posttranslational modification, protein turnover, chaperones; E, Amino acid transport and metabolism; I, Lipid transport and metabolism; M, Cell wall/membrane/envelope biogenesis; P, Inorganic ion transport and metabolism; N, Cell motility; K, Transcription; H, Coenzyme transport and metabolism; V, Defense mechanisms; U, Intracellular trafficking, secretion, and vesicular transport; W, Extracellular structures; D, Cell cycle control, cell division, chromosome partitioning.

The function ‘inorganic ion transport and metabolism’ was mostly represented by proteins involved in iron uptake. Iron is a key element in metabolism, and its availability influences the composition of gut microbiota (Yang et al., 2020). Therefore, iron uptake might confer an advantage to Bacteroidetes growth and survival.

A large proportion of the protein intensity (68-80%) associated with ‘intracellular trafficking, secretion and vesicular transport’ was related to biopolymer transport (ExbB and ExbD). The ExbB and ExbD membrane proteins, along with a third one, form the TonB system that facilitates active transport of specific substances (in this case, biopolymers) across the membrane (Simon et al., 2007). In recent years, biopolymers have gained attention due to their applications in different industries and their role in bacterial pathogenicity. Biopolymers allow bacteria to grow under unfavorable conditions, because they can provide protection, energy storage, or biofilm components (Moradali and Rehm, 2020).

In the functional category of ‘extracellular structures’, all proteins were related to pilus assembly. In S1, FimD/PapC was the most abundant representative protein, and in the other samples, PilF was the most abundant protein. PilF is an ATPase that contributes to the extension of type IV pili. These structures play important roles in pathogenesis by facilitating biofilm formation, adhesion, and motility (Gold et al., 2015). All the above mentioned functions could allow Bacteroidetes to overcome disadvantageous conditions in the human gut and favor its establishment as one of the most abundant phyla.

In contrast to Bacteroidetes, in the Actinobacteria phylum, we identified only a few proteins associated with different functions, mainly ‘carbohydrate transport and metabolism’, ‘translation, ribosomal structure and biogenesis’ and ‘amino acid transport and metabolism’ (Figure 5). Nearly 50% of the proteins in this phylum were associated with ‘carbohydrate transport and metabolism’ consistent with previous studies (Kolmeder et al., 2012). This finding suggests that the members of the Actinobacteria phylum play roles in the degradation of carbohydrates that resist digestion (Pokusaeva et al., 2011).

In the Proteobacteria phylum, most proteins were associated with the functional category ‘cell wall/membrane/envelope biogenesis’. We detected three types of proteins related to this category: (i) ‘opacity proteins and related surface antigens’, (ii) ‘outer membrane protein (porin)’ and (iii) ‘outer membrane protein OmpA and related peptidoglycan-associated lipoproteins’. These proteins are located in the outer membrane and have pathogenic functions; they act as virulence factors and are very immunogenic (Avidan et al., 2008). In S6, opacity proteins were associated with *Enterobacteriaceae*. Porins, which are associated with *Sutterella*, were identified in all samples. OmpA was highly represented in S3, where it comprised nearly 65 % of the protein intensity associated with the ‘cell wall/membrane/envelope biogenesis’ functional category and it was attributed to the species *Bilophila wadsworthia*. When dysbiosis occurs, the Proteobacteria phylum tends to establish itself as the main phylum in the gut community, due in part to its broad adaptability. Further study on the role of Proteobacteria in different health stages might allow us to prevent its spread and avoid the associated health problems.

It is remarkable that, when analyzing each sample individually, we found that some functions were linked to only a few or even a single species. For example, in S3, we observed a strong link between proteins related to ‘coenzyme transport and metabolism’ and the *Prevotella* genus. Of note, this function displayed a protein intensity 2 to 5 times greater in S3 than in the other samples (Table S2). Indeed, over 80% of this COG category was represented by *Prevotella* proteins (Figure 6A). The most abundant species in S3, was *P copri*, which was the principal species related to this COG functional category. Moreover, 80% of the protein intensity associated to ‘coenzyme transport and metabolism’ was related to cobalamin (vitamin B12) metabolism. The behavior of *P. copri* largely depends on the strain (De Filippis et al., 2019). Strains from Western individuals were associated with the synthesis of different B vitamins (De Filippis et al., 2019), consistent with our results. Most human gut taxa require B12, but the majority of these taxa lack de novo B12 synthesis; therefore most of these bacteria rely on cobalamin-uptake mechanisms to acquire sufficient B12 (Degnan et al., 2014a). It has been hypothesized that microbial communities might be manipulated to promote health by changing vitamin intake, due to the high competition for cobalamin (Degnan et al., 2014b). Members of Firmicutes and Actinobacteria phyla harbor complete B12 biosynthetic pathways (Rowley and Kendall, 2019). In contrast, Bacteroidetes, which lacks de novo B12 biosynthetic genes, encodes several cobalamin transporters (Degnan et al., 2014a), which could be detected among the identified proteins. Interestingly, the cobalamin biosynthesis protein, Cbik (COG4822), which is one of the first proteins that participates in de novo cobalamin synthesis, was related to Firmicutes. In contrast, an outer membrane cobalamin receptor protein (COG4206) was associated with *Prevotella*. Aditionally some proteins required for the activation of B12 synthesis were associated with *Bacteroides* (e.g., cobalamin biosynthesis protein CobN; COG1429) and *Prevotella* (e.g., cobalamin adenosyltransferase; COG2096). Furthermore, vitamin B12 consumption has been related to an increase in the relative abundance of *Prevotella* over *Bacteroides* (Carrothers et al., 2015), which might potentially explain the high abundance of the *Prevotella* genus observed in S3.

**Figure 6.**
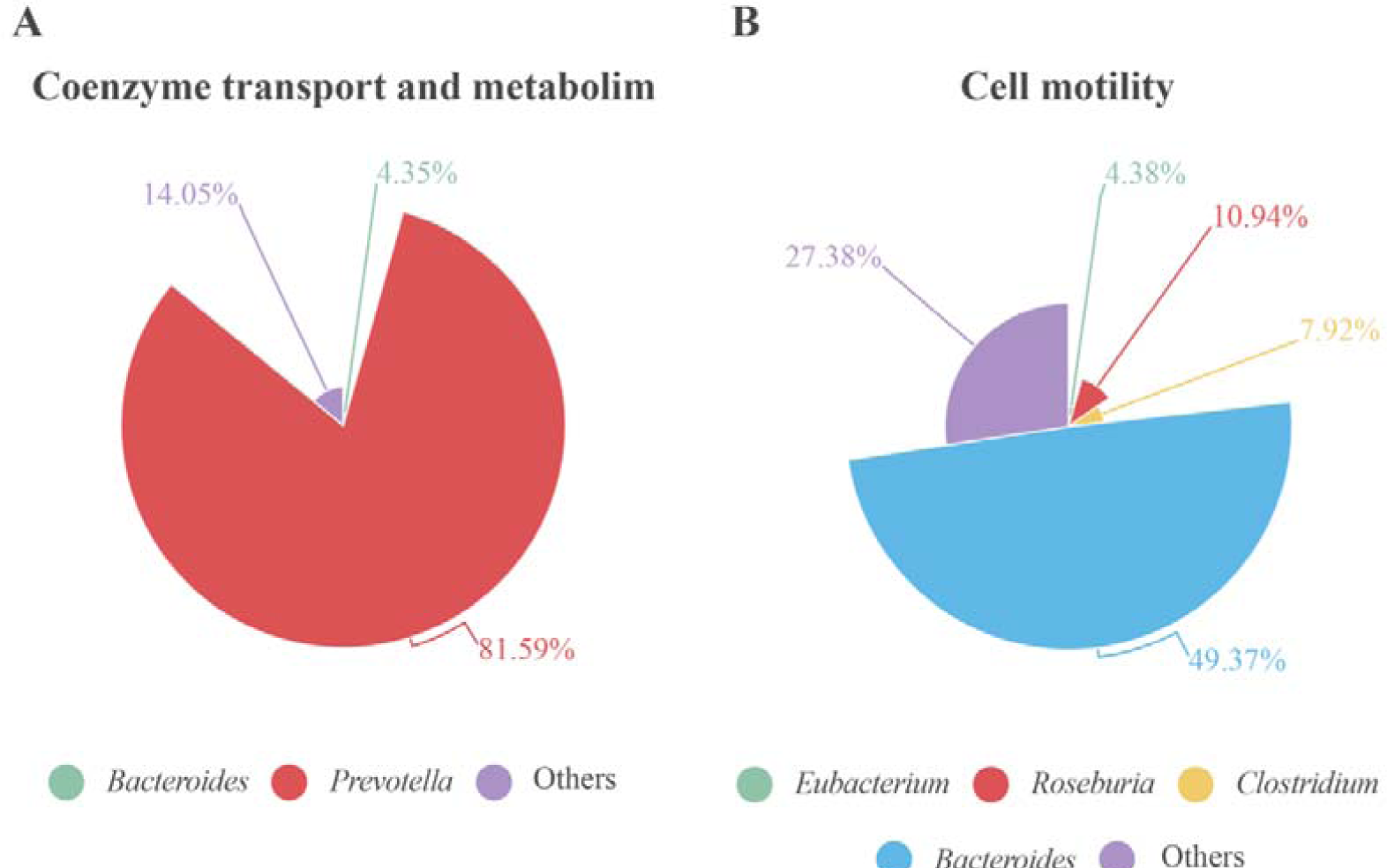
Proteins associated with (A) coenzyme transport and metabolism in S3 and (B) cell motility in S5 at genus level. Contribution of each genus is represented as the percentage of protein intensity of the genus assigned to each COG category.

Another example of this taxa-function link was observed in S5. We found that the intensity and the abundance of the functional category ‘cell motility’ was higher in S5 than in the other samples. In this functional category, Firmicutes and Bacteroidetes proteins represented nearly 50% of the protein intensity. Within the Bacteroidetes phylum, *Bacteroides* was the only genus whose proteins contributed to the cell motility category in S5(Figure 6B). We found that proteins from *B. vulgatus* were responsible for 33% of the total intensity for this function. Furthermore, all proteins associated with the *Bacteroides* genus that were related to the ‘cell motility’ category belonged to the ‘flagellar hook-associated protein FlgK’ group. In contrast, Firmicutes contributed more diverse taxa. The principal genera were *Roseburia* (11%), *Clostridium* (8%), and *Eubacterium* (4%), which all belong to the Clostridiales order (Figure 6B). These genera are also associated with flagellin proteins (Kolmeder et al., 2012). Similar to *Bacteroides* genus, all the Firmicutes proteins involved in the ‘cell motility’ category belonged to the ‘flagellin and related hook-associated protein FlgL’ group. These two flagellar proteins (Flgk and FlgL) were among the 30 most abundant proteins. Flgk associates with FlgL to form the junction between the flagellin filament and the hook in the flagellum (Hong et al., 2018). The flagellum serves as a virulence factor, because it contributes to colonization and the invasion of host surfaces (Hong et al., 2018). Flagellin can also trigger host immune responses through Toll-like receptors (Ramos et al., 2004). In addition, flagellin facilitates bacterial access to food sources. Taken together, these findings suggested that flagellar proteins are key elements for these bacteria to survive in the gut.

These two examples are consistent with the premise that, in each individual, different biological functions are linked to specific taxa that contribute to specific host-microbiota interactions. This principle should be keep in mind, particularly in the field of personalized medicine and nutrition (Mills et al., 2019).

Another important function of the gut microbiota in human physiology is the synthesis of short chain fatty acids (SCFA) through the fermentation of different non-digestible carbohydrates. SCFAs are recognized by G-coupled receptors that trigger the secretion of intestinal peptides (Cani, 2018). The main SCFAs are butyrate, propionate, and acetate. Each of these SCFAs plays important roles in human health, such as regulating the epithelial barrier integrity, providing anti-inflammatory effects, and serving as the primary nutrient for colonocytes (Morrison and Preston, 2016; Yamamura et al., 2020). Butyrate is the most important energy source for colonocytes. Moreover, it influences the microbial environment and ecology, and it prevents the expansion of pathogens (Cani, 2018). Due to current interest in the beneficial roles that SCFAs perform in the intestine (Riviere et al., 2016), we analyzed SCFA production by Firmicutes in all 6 samples. Nineteen KOs were associated with the KEGG pathway involved in ‘butanoate metabolism’ (Table S4). Firmicutes proteins represented 62% of the total butyrate pathway protein intensity, and the main genus was *Faecalibacterium* (a member of *Ruminococcaceae)*, which comprised 25% of the intensity. These results were consistent with previous studies that showed that members of the Firmicutes phylum were the main butyrate producers, which highlighted the role of *Ruminococcaceae* family in butyrate production (Louis et al., 2009; Morrison and Preston, 2016; Yamamura et al., 2020). The KOs identified in butyrate production from acetyl-CoA were all assigned to the Firmicutes phylum. The KOs related to SCFA production that were assigned to Bacteroidetes were not directly involved in the biosynthetic pathway, instead, they were related to acetyl-CoA production from pyruvate and the interconversion of succinate-fumarate (Figure S1).

#### 3.3.3 Human proteins detected in proteomic studies of stool samples

The Metalab program allows the identification of human proteins that are present in stool samples. We identified up to 92 human proteins throughout the 6 human samples. Interestingly, although human peptides only accounted for 2% of the total identified peptides, the intensities of these peptides represented up to 13% of the total peptide intensity in S1. This difference between the abundance of identified peptides and their intensity was consistent with findings in a previous study (Zhang et al., 2017). The most abundant proteins were chymotrypsin C, trypsin 2, phospholipase A2, and alpha amylase. Alpha amylase, a starch degradation protein, is present in human epithelial cells, where it induces cell proliferation and differentiation (Date et al., 2020).

A group of 35 human proteins that were found in all stool samples at high abundance (Figure 7 and Table S5) were selected for STRING analysis. These proteins represented over 80% of the total intensity of identified human proteins. Most of these selected proteins were also found in previous studies on the microbiome (Zhang et al., 2017; Zhong et al., 2019), including intestinal mucin proteins (MUC2, MUC5B, and MUC13) and digestive enzymes, like chymotrypsinogen B2 and carboxypeptidases. MUC2 is the predominant mucin type in the stomach mucus layer. Intestinal mucus provides a protective, lubricating barrier against particles and infectious agents, and it interacts with microorganisms. For example, microbiota can feed on mucin glycans and convert them into SCFAs that supply colonocytes and other gut epithelial cells with energy (Ouwerkerk et al., 2013). Interestingly, MUC13 is abundant in many adenocarcinomas (Sheng et al., 2019), where it serves as a potential prognostic factor (Filippou et al., 2018).

**Figure 7.**
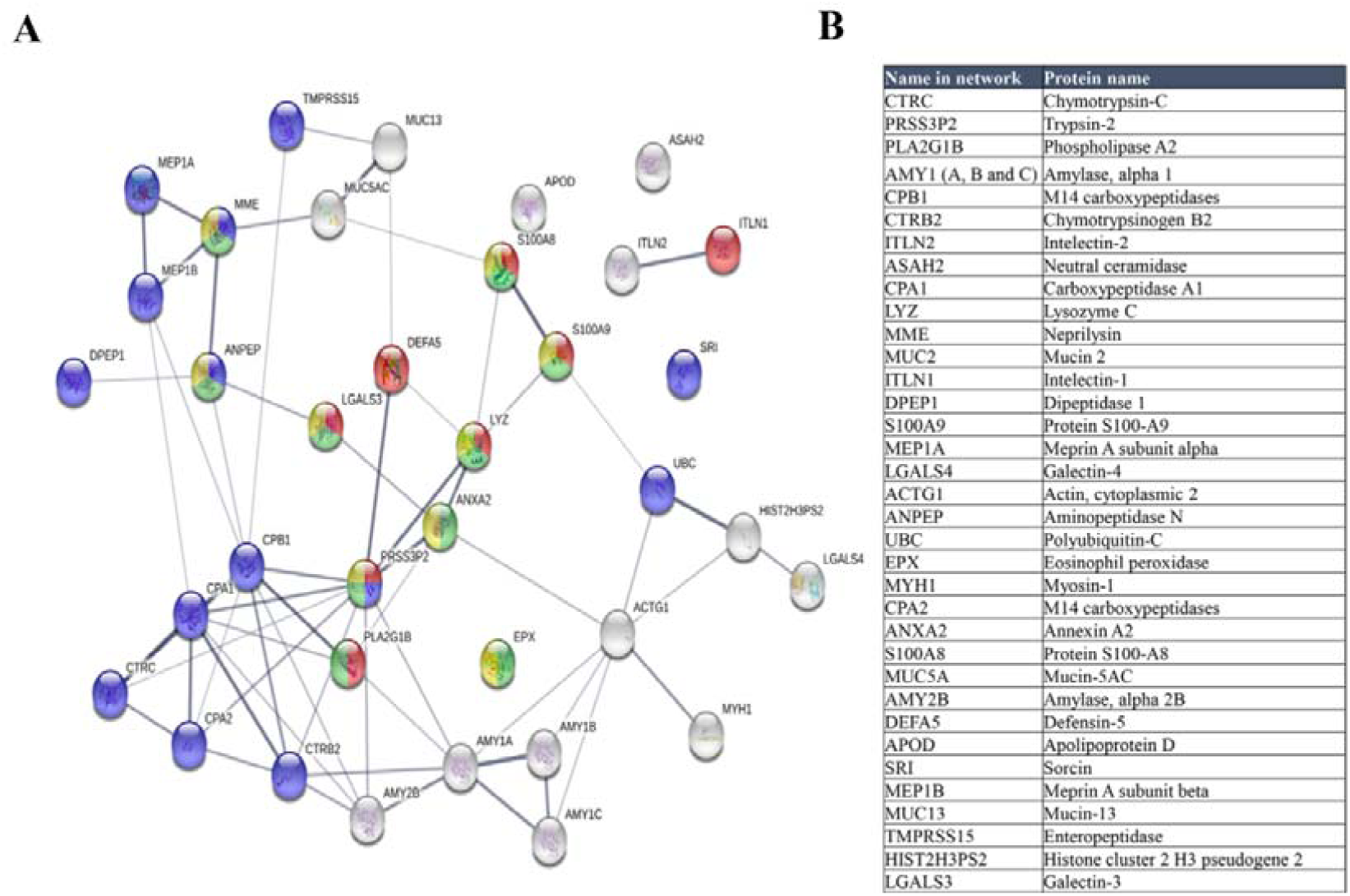
(A) STRING map protein-protein interaction network of the 35 more abundant human proteins identified in this study. Line thickness indicates the strength of data support. The colours represent different processes; red: antimicrobial humoral response, green: neutrophil mediated immunity, blue: proteolysis and yellow: neutrophil degranulation. (B)Table of these proteins ordered according to decreasing abundance.

When we analyzed the selected 35 human proteins with the STRING program (version 11.0), only one protein (Mucin 2) did not match in the STRING database. The other proteins, showed significant protein-protein interactions (P=1.0 ×10^−16^, Figure 7). The antimicrobial humoral response was the most significant enrichment process (P=3.10 ×10^−07^), followed by proteolysis (P=7.32 ×10^−06^), and immune system-related processes, including neutrophil-mediated immunity (P= 7.32 ×10^−06^), neutrophil degranulation (P= 3.35 ×10^−05^), granulocyte migration (P=3.4 ×10^−05^), and immune effector process (P=4.84 ×10^−05^).

We found that the enrichment of the antimicrobial humoral response process was due to the high number of antimicrobial peptides identified in the samples, consistent with other metaproteomics studies (Zhong et al., 2019). These antimicrobial peptides, like defensin-5, lysozyme c, and phospholipase A2, accomplish important roles in the defense against bacteria (Zhong et al., 2019). Apart from antimicrobial peptides, we identified other proteins like Interlectin 1, S100A9, S100A8, and galectin 3, which are related to the antimicrobial humoral response. Interlectin 1 is a lectin that binds microbial glycans and is used by the immune system to discriminate human cells from microbes (Wesener et al., 2015). S100A9 and S100A8 can induce neutrophil chemotaxis and adhesion, and they are found as a complex (S100A8/A9) called calprotectin. High levels of calprotectin have been observed in inflammatory bowel disease, and thus, it was proposed as a marker (Fukunaga et al., 2018). Galectin 3 is a lectin involved in neutrophil activation and adhesion. Galectin 3 plays an important role in inflammation and is associated with several diseases (Sciacchitano et al., 2018).

Proteolysis was also highly represented in the STRING results. Proteases play important roles in gastrointestinal disorders (Antalis et al., 2007). Enteropeptidase is responsible for activating the conversion of pancreatic trypsinogen to trypsin, which activates other proenzymes (e.g., chymotrypsinogen, procarboxypeptidases, and others). Trypsin is also involved in processing defensins. The most significantly enriched functions were metallopeptidase activity, hydrolase activity, and peptidase activity (P=5.23 ×10^−09^). We identified the metalloprotease, meprin A (alpha and beta subunits), which is implicated in inflammation and tissue remodeling (Kaushal et al., 2013).

We also observed a cellular component enrichment. Proteins related to the extracellular space (P=3.59 ×10^−19^) and extracellular regions (P=5.40 ×10^−18^) were the most significantly enriched. This finding was consistent with the fact that these extracellular proteins might have been dragged through the human intestine during fecal sampling.

## 4 Concluding remarks

In this study, we designed and evaluated different stool sample processing methods for the metaproteomic study of gut microbiota. We found that proper moistening of the fecal sample prior to protein extraction was critical for good protein recovery and peptide identification. Furthermore, the cell breaking method determined the number of peptides that could be identified and, more interestingly, the particular taxa that would be enriched in the metaproteomic taxonomic profile. An ultrasonication procedure prior to bead-beating in microbial cell disruption raised the numbers of proteins and peptides identified, and thus, the number of taxa identified. However, bead-beating alone increased the number of Proteobacteria proteins identified. This method could be more informative for studies related to diseases, because Proteobacteria phylum was associated with different dysbiosis stages, which facilitated the differentiation of healthy and unhealthy stages. These results are relevant to future metaproteomic studies on the human gut microbiota, because they inform the selection of the best protocol, based on the specific interest in a particular taxon-related disease. It is worth mentioning that we could identify various taxa, including Bacteria, Archaea, and eukaryotic microbial cells, like Fungi.

The metaproteomic studies enabled the profiling of protein functions in the microbiota from 6 healthy individuals. We found interesting correlations between specific microbial functions relevant to the host and the main taxa involved. For example, vitamin B12 process was mainly produced by proteins in the *Prevotella* genus. This could be particularly interesting when searching for links between certain taxon-related proteins and specific beneficial or detrimental host stages in future health-disease metaproteomic studies. Finally, we also detected 92 human proteins, which were mostly of them related to the antimicrobial humoral response.

The newly described protocols for enriching specific taxa can facilitate the functional analysis of both microbial and human proteins in the human gut. This information can improve the design of future metaproteomic studies on gut microbiota and open up new prospects in the field of host-pathogen interactions in different diseases.

## Supplementary material

**Table S1. Taxonomy of human gut microbiota of 6 healthy adult individuals**.

**Table S2. Functional analysis of human gut microbiota of 6 healthy adults. The abundance of each COG category is represented as the sum intensity of proteins annotated to the COG category in each sample. The box color indicates the grade of intensity within the 6 samples. Intensities with the number in red are those of special interest and discussed in deep in the text**.

**Table S3. Thirty most abundant microbial proteins found in the study**.

**Table S4. KOs entries belonging to butanoate metabolism pathway found in the study**.

**Table S5. Abundant human proteins used for STRING analysis**.

**Figure S1.**
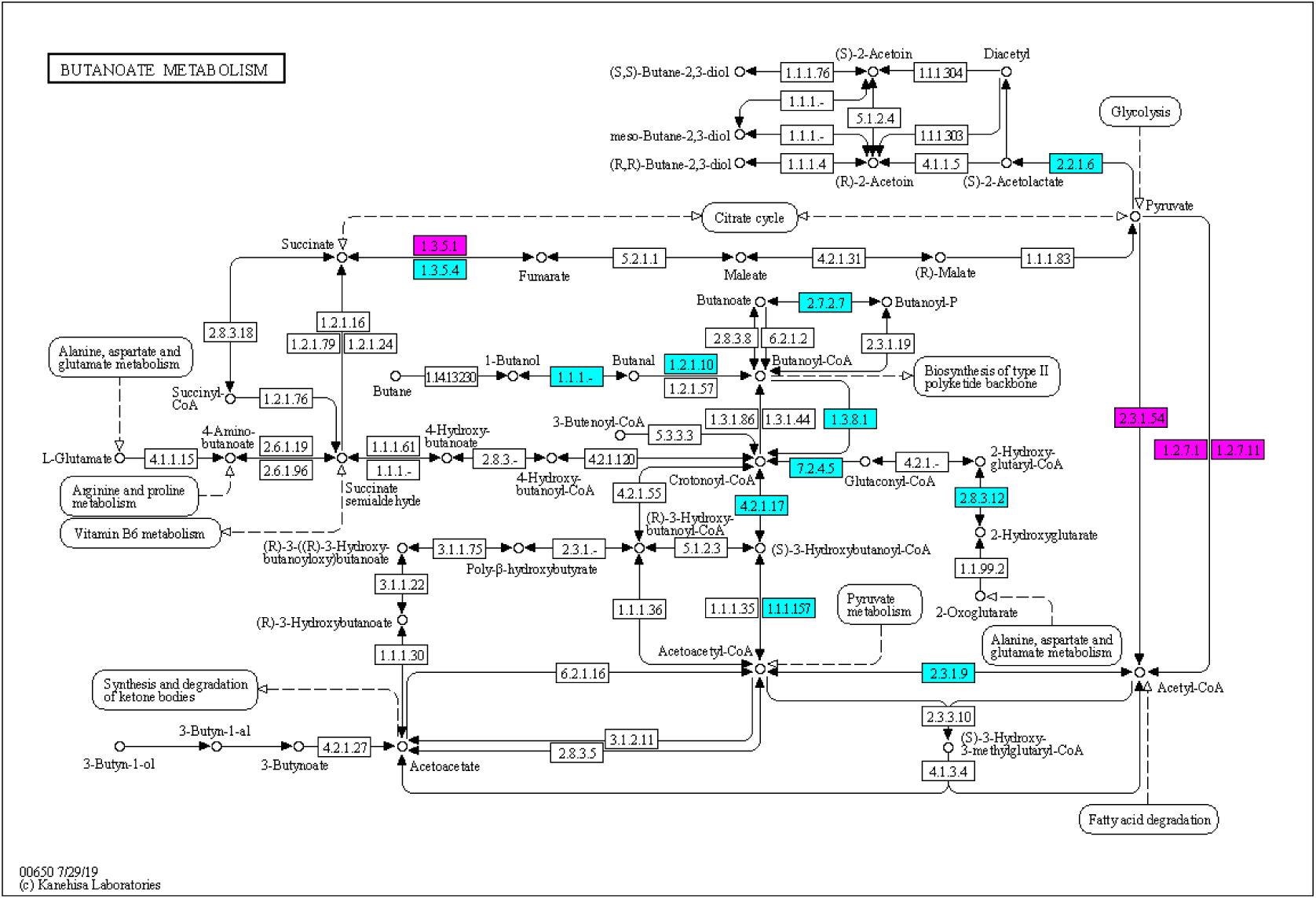
Butyrate metabolism KEGG pathway downloaded from KEGG website (**http://www.kegg.jp**). Coloured KO numbers indicate proteins identified in this study. Firmicutes proteins are in coloured in blue and Bacteroidetes proteins in purple.

## Conflict of Interest

The authors declare that the research was conducted in the absence of any commercial or financial relationships that could be construed as a potential conflict of interest.

## Author contributions

C.G-D, R.M-L and I.Z performed the experiments.

C.G-D analysed, interpreted the data, and wrote the manuscript.

R.M-L and A.P contribute to the data analysis and to the writing of the manuscript.

A.P carried out the statistical analysis.

E.P and E.R supported the bioinformatic analysis.

M.L. designed and supervised the proteomic experiments.

J.A. participated in data analyses and critically reviewed the manuscript.

C.G. and L.M. conceived and designed the experiments, supervised the experimental work and critically reviewed the manuscript.

All the authors approved the final version of the manuscript.

## Acknowledgements

We thank Dr. Figeys, Dr. Zhibin Ning and Dr. Xu Zang for their help with the MetaLab software, Pedro Botías and Dr. Jesús García-Cantalejo (Genomics Unit, UCM) for helpful discussions and VidaCord for the financial support. This study was supported by InGEMICS-CM B2017/BMD3691 from the Comunidad de Madrid, RTI2018-094004-B-100 from Spanish Ministry of Science and Innovation, Spanish Network for the Research in Infectious Diseases (REIPI RD16/0016/0011) and PRB3 (PT17/0019/0012) from the ISCIII. InGEMICSCM, REIPI and PRB3 are cofinanced by European Development Regional Fund ERDF “A way to achieve Europe”. These results are lined up with the Human Infectious Diseases HPP initiative from the Human Proteome Project (HID-HPP). The proteomics analyses were performed in the Proteomics facility of Complutense University of Madrid (UCM) a member of ProteoRed-ISCIII network. Carmen García received a grant from the European Social Fund in the Operational Programme of Youth Employment, and from the Youth Employment Initiative (YEI) fund by the Comunidad de Madrid (PEJD-2019-PRE/BMD-16854).

## Notes

http://www.proteomexchange.org

### Competing Interest Statement

The authors have declared no competing interest.

## Bibliography

Adak, A., and Khan, M.R. (2019). An insight into gut microbiota and its functionalities. Cell. Mol. Life Sci. 76(3), 473–493. doi: 10.1007/s00018-018-2943-4.

Alarcón, T., D’Auria, G., Delgado, S., Del Campo, R., and Ferrer, M. (2016). “Microbiota,” in Procedimientos en Microbiología Clínica, eds. E. Cercenado & R. Cantón. 2 ed (Madrid: Sociedad Española de Enfermedades Infecciosas y Microbiología Clínica (SEIMC)), 1–6.

Antalis, T.M., Shea-Donohue, T., Vogel, S.N., Sears, C., and Fasano, A. (2007). Mechanisms of disease: protease functions in intestinal mucosal pathobiology. Nat. Clin. Pract. Gastroenterol. Hepatol. 4(7), 393–402. doi: 10.1038/ncpgasthep0846.

Arumugam, M., Raes, J., Pelletier, E., Le Paslier, D., Yamada, T., Mende, D.R., et al. (2011). Enterotypes of the human gut microbiome. Nature 473(7346), 174–180. doi: 10.1038/nature09944.

Avidan, O., Kaltageser, E., Pechatnikov, I., Wexler, H.M., Shainskaya, A., and Nitzan, Y. (2008). Isolation and characterization of porins from Desulfovibrio piger and Bilophila wadsworthia: structure and gene sequencing. Arch. Microbiol. 190(6), 641. doi: 10.1007/s00203-008-0416-0.

Bjørkhaug, S.T., Aanes, H., Neupane, S.P., Bramness, J.G., Malvik, S., Henriksen, C., et al. (2019). Characterization of gut microbiota composition and functions in patients with chronic alcohol overconsumption. Gut Microbes 10(6), 663–675. doi: 10.1080/19490976.2019.1580097.

Blackburn, J.M., and Martens, L. (2016). The challenge of metaproteomic analysis in human samples. Expert Rev. Proteomics 13(2), 135–138. doi: 10.1586/14789450.2016.1135058.

Cani, P.D. (2018). Human gut microbiome: hopes, threats and promises. Gut 67(9), 1716–1725. doi: 10.1136/gutjnl-2018-316723.

Carrothers, J.M., York, M.A., Brooker, S.L., Lackey, K.A., Williams, J.E., Shafii, B., et al. (2015). Fecal Microbial Community Structure Is Stable over Time and Related to Variation in Macronutrient and Micronutrient Intakes in Lactating Women. J. Nutr. 145(10), 2379–2388. doi: 10.3945/jn.115.211110.

Cheng, K., Ning, Z., Zhang, X., Li, L., Liao, B., Mayne, J., et al. (2017). MetaLab: an automated pipeline for metaproteomic data analysis. Microbiome 5(1), 157. doi: 10.1186/s40168-017-0375-2.

Consortium, U. (2019). UniProt: a worldwide hub of protein knowledge. Nucleic Acids Res. 47(D1), D506–d515. doi: 10.1093/nar/gky1049.

Date, K., Yamazaki, T., Toyoda, Y., Hoshi, K., and Ogawa, H. (2020). α-Amylase expressed in human small intestinal epithelial cells is essential for cell proliferation and differentiation. J. Cell Biochem. 121(2), 1238–1249. doi: 10.1002/jcb.29357.

David, L.A., Maurice, C.F., Carmody, R.N., Gootenberg, D.B., Button, J.E., Wolfe, B.E., et al. (2014). Diet rapidly and reproducibly alters the human gut microbiome. Nature 505(7484), 559–563. doi: 10.1038/nature12820.

De Filippis, F., Pasolli, E., Tett, A., Tarallo, S., Naccarati, A., De Angelis, M., et al. (2019). Distinct Genetic and Functional Traits of Human Intestinal Prevotella copri Strains Are Associated with Different Habitual Diets. Cell Host Microbe 25(3), 444–453.e443. doi: 10.1016/j.chom.2019.01.004.

Debyser, G., Mesuere, B., Clement, L., Van de Weygaert, J., Van Hecke, P., Duytschaever, G., et al. (2016). Faecal proteomics: A tool to investigate dysbiosis and inflammation in patients with cystic fibrosis. J. Cyst. Fibros. 15(2), 242–250. doi: 10.1016/j.jcf.2015.08.003.

Degnan, P.H., Barry, N.A., Mok, K.C., Taga, M.E., and Goodman, A.L. (2014a). Human gut microbes use multiple transporters to distinguish vitamin B12 analogs and compete in the gut. Cell Host Microbe 15(1), 47–57. doi: 10.1016/j.chom.2013.12.007.

Degnan, P.H., Taga, M.E., and Goodman, A.L. (2014b). Vitamin B12 as a modulator of gut microbial ecology. Cell Metab. 20(5), 769–778. doi: 10.1016/j.cmet.2014.10.002.

Dridi, B., Henry, M., El Khéchine, A., Raoult, D., and Drancourt, M. (2009). High Prevalence of Methanobrevibacter smithii and Methanosphaera stadtmanae Detected in the Human Gut Using an Improved DNA Detection Protocol. PLoS One 4(9), e7063. doi: 10.1371/journal.pone.0007063.

Everard, A., Belzer, C., Geurts, L., Ouwerkerk, J.P., Druart, C., Bindels, L.B., et al. (2013). Cross-talk between Akkermansia muciniphila and intestinal epithelium controls diet-induced obesity. Proc. Natl. Acad. Sci. U. S. A. 110(22), 9066–9071. doi: 10.1073/pnas.1219451110.

Fattorusso, A., Di Genova, L., Dell’Isola, G.B., Mencaroni, E., and Esposito, S. (2019). Autism Spectrum Disorders and the Gut Microbiota. Nutrients 11(3). doi: 10.3390/nu11030521.

Filippou, P.S., Ren, A.H., Korbakis, D., Dimitrakopoulos, L., Soosaipillai, A., Barak, V., et al. (2018). Exploring the potential of mucin 13 (MUC13) as a biomarker for carcinomas and other diseases. Clin. Chem. Lab. Med. 56(11), 1945–1953. doi: 10.1515/cclm-2018-0139.

Franke, T., and Deppenmeier, U. (2018). Physiology and central carbon metabolism of the gut bacterium Prevotella copri. Mol. Microbiol. 109(4), 528–540. doi: 10.1111/mmi.14058.

Fukunaga, S., Kuwaki, K., Mitsuyama, K., Takedatsu, H., Yoshioka, S., Yamasaki, H., et al. (2018). Detection of calprotectin in inflammatory bowel disease: Fecal and serum levels and immunohistochemical localization. Int. J. Mol. Med. 41(1), 107–118. doi: 10.3892/ijmm.2017.3244.

Gold, V.A.M., Salzer, R., Averhoff, B., and Kühlbrandt, W. (2015). Structure of a type IV pilus machinery in the open and closed state. eLife 4, e07380. doi: 10.7554/eLife.07380.

Hiippala, K., Kainulainen, V., Kalliomäki, M., Arkkila, P., and Satokari, R. (2016). Mucosal Prevalence and Interactions with the Epithelium Indicate Commensalism of Sutterella spp. Front. Microbiol. 7, 1706–1706. doi: 10.3389/fmicb.2016.01706.

Hoffmann, C., Dollive, S., Grunberg, S., Chen, J., Li, H., Wu, G.D., et al. (2013). Archaea and fungi of the human gut microbiome: correlations with diet and bacterial residents. PLoS One 8(6), e66019. doi: 10.1371/journal.pone.0066019.

Hong, H.J., Kim, T.H., Song, W.S., Ko, H.-J., Lee, G.-S., Kang, S.G., et al. (2018). Crystal structure of FlgL and its implications for flagellar assembly. Sci. Rep. 8(1), 14307. doi: 10.1038/s41598-018-32460-9.

Issa Isaac, N., Philippe, D., Nicholas, A., Raoult, D., and Eric, C. (2019). Metaproteomics of the human gut microbiota: Challenges and contributions to other OMICS. Clin. Mass Spectrom. 14, 18–30. doi: 10.1016/j.clinms.2019.06.001.

Jakobsson, H.E., Jernberg, C., Andersson, A.F., Sjölund-Karlsson, M., Jansson, J.K., and Engstrand, L. (2010). Short-term antibiotic treatment has differing long-term impacts on the human throat and gut microbiome. PLoS One 5(3), e9836. doi: 10.1371/journal.pone.0009836.

Jandhyala, S.M., Talukdar, R., Subramanyam, C., Vuyyuru, H., Sasikala, M., and Nageshwar Reddy, D. (2015). Role of the normal gut microbiota. World J. Gastroenterol. 21(29), 8787–8803. doi: 10.3748/wjg.v21.i29.8787.

Jin, Y., Wu, S., Zeng, Z., and Fu, Z. (2017). Effects of environmental pollutants on gut microbiota. Environ. Pollut. 222, 1–9. doi: 10.1016/j.envpol.2016.11.045.

Jovel, J., Patterson, J., Wang, W., Hotte, N., O’Keefe, S., Mitchel, T., et al. (2016). Characterization of the Gut Microbiome Using 16S or Shotgun Metagenomics. Front. Microbiol. 7(459). doi: 10.3389/fmicb.2016.00459.

Kaushal, G.P., Haun, R.S., Herzog, C., and Shah, S.V. (2013). Meprin A metalloproteinase and its role in acute kidney injury. Am. J. Physiol. Renal. Physiol. 304(9), F1150–1158. doi: 10.1152/ajprenal.00014.2013.

Kim, S., and Jazwinski, S.M. (2018). The Gut Microbiota and Healthy Aging: A Mini-Review. Gerontology 64(6), 513–520. doi: 10.1159/000490615.

Kolmeder, C.A., de Been, M., Nikkilä, J., Ritamo, I., Mättö, J., Valmu, L., et al. (2012). Comparative Metaproteomics and Diversity Analysis of Human Intestinal Microbiota Testifies for Its Temporal Stability and Expression of Core Functions. PLoS One 7(1), e29913. doi: 10.1371/journal.pone.0029913.

Kolmeder, C.A., Salojarvi, J., Ritari, J., de Been, M., Raes, J., Falony, G., et al. (2016). Faecal Metaproteomic Analysis Reveals a Personalized and Stable Functional Microbiome and Limited Effects of a Probiotic Intervention in Adults. PLoS One 11(4), e0153294. doi: 10.1371/journal.pone.0153294.

Kowalski, K., and Mulak, A. (2019). Brain-Gut-Microbiota Axis in Alzheimer’s Disease. J. Neurogastroenterol. Motil. 25(1), 48–60. doi: 10.5056/jnm18087.

La Reau, A.J., Meier-Kolthoff, J.P., and Suen, G. (2016). Sequence-based analysis of the genus Ruminococcus resolves its phylogeny and reveals strong host association. Microb. Genom. 2(12), e000099. doi: 10.1099/mgen.0.000099.

Li, J., Jia, H., Cai, X., Zhong, H., Feng, Q., Sunagawa, S., et al. (2014). An integrated catalog of reference genes in the human gut microbiome. Nat. Biotechnol. 32(8), 834–841. doi: 10.1038/nbt.2942.

Lin, D., Peters, B.A., Friedlander, C., Freiman, H.J., Goedert, J.J., Sinha, R., et al. (2018). Association of dietary fibre intake and gut microbiota in adults. Br. J. Nutr. 120(9), 1014–1022. doi: 10.1017/s0007114518002465.

Lloyd-Price, J., Abu-Ali, G., and Huttenhower, C. (2016). The healthy human microbiome. Genome Med. 8(1), 51–51. doi: 10.1186/s13073-016-0307-y.

Louis, P., Young, P., Holtrop, G., and Flint, H. (2009). Diversity of human colonic butyrate-producing bacteria revealed by analysis of the butyryl-CoA:acetate CoA-transferase gene. Environ. Microbiol. 12, 304–314. doi: 10.1111/j.1462-2920.2009.02066.x.

Mills, S., Stanton, C., Lane, J.A., Smith, G.J., and Ross, R.P. (2019). Precision Nutrition and the Microbiome, Part I: Current State of the Science. Nutrients 11(4), 923. doi: 10.3390/nu11040923.

Moradali, M.F., and Rehm, B.H.A. (2020). Bacterial biopolymers: from pathogenesis to advanced materials. Nat. Rev. Microbiol. 18(4), 195–210. doi: 10.1038/s41579-019-0313-3.

Morrison, D.J., and Preston, T. (2016). Formation of short chain fatty acids by the gut microbiota and their impact on human metabolism. Gut Microbes 7(3), 189–200. doi: 10.1080/19490976.2015.1134082.

Natividad, J.M., Lamas, B., Pham, H.P., Michel, M.-L., Rainteau, D., Bridonneau, C., et al. (2018). Bilophila wadsworthia aggravates high fat diet induced metabolic dysfunctions in mice. Nat. Commun. 9(1), 2802–2802. doi: 10.1038/s41467-018-05249-7.

Ni, J., Wu, G.D., Albenberg, L., and Tomov, V.T. (2017). Gut microbiota and IBD: causation or correlation? Nat. Rev. Gastroenterol. Hepatol. 14(10), 573–584. doi: 10.1038/nrgastro.2017.88.

Ouwerkerk, J.P., de Vos, W.M., and Belzer, C. (2013). Glycobiome: bacteria and mucus at the epithelial interface. Best. Pract. Res. Clin. Gastroenterol. 27(1), 25–38. doi: 10.1016/j.bpg.2013.03.001.

Pitarch, A., Nombela, C., and Gil, C. (2008). Cell Wall Fractionation for Yeast and Fungal Proteomics. Methods Mol. Biol. 425, 217–239. doi: 10.1007/978-1-60327-210-0_19.

Pitarch, A., Nombela, C., and Gil, C. (2009). Proteomic profiling of serologic response to Candida albicans during host-commensal and host-pathogen interactions. Methods Mol. Biol. 470, 369–411. doi: 10.1007/978-1-59745-204-5_26.

Pokusaeva, K., Fitzgerald, G.F., and van Sinderen, D. (2011). Carbohydrate metabolism in Bifidobacteria. Genes Nutr. 6(3), 285–306. doi: 10.1007/s12263-010-0206-6.

Ramos, H.C., Rumbo, M., and Sirard, J.C. (2004). Bacterial flagellins: mediators of pathogenicity and host immune responses in mucosa. Trends Microbiol. 12(11), 509–517. doi: 10.1016/j.tim.2004.09.002.

Riviere, A., Selak, M., Lantin, D., Leroy, F., and De Vuyst, L. (2016). Bifidobacteria and Butyrate-Producing Colon Bacteria: Importance and Strategies for Their Stimulation in the Human Gut. Front. Microbiol. 7, 979. doi: 10.3389/fmicb.2016.00979.

Rizzatti, G., Lopetuso, L.R., Gibiino, G., Binda, C., and Gasbarrini, A. (2017). Proteobacteria: A Common Factor in Human Diseases. BioMed Res. Int. 2017, 9351507–9351507. doi: 10.1155/2017/9351507.

Rowley, C.A., and Kendall, M.M. (2019). To B12 or not to B12: Five questions on the role of cobalamin in host-microbial interactions. PLoS Pathog. 15(1), e1007479. doi: 10.1371/journal.ppat.1007479.

Saffouri, G.B., Shields-Cutler, R.R., Chen, J., Yang, Y., Lekatz, H.R., Hale, V.L., et al. (2019). Small intestinal microbial dysbiosis underlies symptoms associated with functional gastrointestinal disorders. Nat. Commun. 10(1), 2012. doi: 10.1038/s41467-019-09964-7.

Salonen, A., Nikkilä, J., Jalanka-Tuovinen, J., Immonen, O., Rajilić-Stojanović, M., Kekkonen, R.A., et al. (2010). Comparative analysis of fecal DNA extraction methods with phylogenetic microarray: effective recovery of bacterial and archaeal DNA using mechanical cell lysis. J. Microbiol. Methods 81(2), 127–134. doi: 10.1016/j.mimet.2010.02.007.

Scher, J.U., Sczesnak, A., Longman, R.S., Segata, N., Ubeda, C., Bielski, C., et al. (2013). Expansion of intestinal Prevotella copri correlates with enhanced susceptibility to arthritis. eLife 2, e01202–e01202. doi: 10.7554/eLife.01202.

Sciacchitano, S., Lavra, L., Morgante, A., Ulivieri, A., Magi, F., De Francesco, G.P., et al. (2018). Galectin-3: One Molecule for an Alphabet of Diseases, from A to Z. Int. J. Mol. Sci. 19(2). doi: 10.3390/ijms19020379.

Sheng, Y.H., Wong, K.Y., Seim, I., Wang, R., He, Y., Wu, A., et al. (2019). MUC13 promotes the development of colitis-associated colorectal tumors via β-catenin activity. Oncogene 38(48), 7294–7310. doi: 10.1038/s41388-019-0951-y.

Shin, N.R., Whon, T.W., and Bae, J.W. (2015). Proteobacteria: microbial signature of dysbiosis in gut microbiota. Trends Biotechnol. 33(9), 496–503. doi: 10.1016/j.tibtech.2015.06.011.

Simon, M.I., Crane, B., and Crane, A. (2007). Two-Component Signaling Systems, Part B. San Diego, United States: Elsevier Science Publishing Co Inc.

Tanca, A., Abbondio, M., Palomba, A., Fraumene, C., Manghina, V., Cucca, F., et al. (2017). Potential and active functions in the gut microbiota of a healthy human cohort. Microbiome 5(1). doi: 10.1186/s40168-017-0293-3.

Tanca, A., Palomba, A., Pisanu, S., Addis, M.F., and Uzzau, S. (2015). Enrichment or depletion? The impact of stool pretreatment on metaproteomic characterization of the human gut microbiota. Proteomics 15(20), 3474–3485. doi: 10.1002/pmic.201400573.

Tett, A., Huang, K.D., Asnicar, F., Fehlner-Peach, H., Pasolli, E., Karcher, N., et al. (2019). The Prevotella copri Complex Comprises Four Distinct Clades Underrepresented in Westernized Populations. Cell Host Microbe 26(5), 666–679.e667. doi: 10.1016/j.chom.2019.08.018.

Wang, H.-X., and Wang, Y.-P. (2016). Gut Microbiota-brain Axis. Chin. Med. J. 129(19), 2373–2380. doi: 10.4103/0366-6999.190667.

Wang, Z., Wang, Q., Zhao, J., Gong, L., Zhang, Y., Wang, X., et al. (2019). Altered diversity and composition of the gut microbiome in patients with cervical cancer. AMB Express 9(1), 40. doi: 10.1186/s13568-019-0763-z.

Wesener, D.A., Wangkanont, K., McBride, R., Song, X., Kraft, M.B., Hodges, H.L., et al. (2015). Recognition of microbial glycans by human intelectin-1. Nat. Struct. Mol. Biol. 22(8), 603–610. doi: 10.1038/nsmb.3053.

Wu, G.D., Chen, J., Hoffmann, C., Bittinger, K., Chen, Y.-Y., Keilbaugh, S.A., et al. (2011). Linking long-term dietary patterns with gut microbial enterotypes. Science (New York, N.Y.) 334(6052), 105–108. doi: 10.1126/science.1208344.

Yamamura, R., Nakamura, K., Kitada, N., Aizawa, T., Shimizu, Y., Nakamura, K., et al. (2020). Associations of gut microbiota, dietary intake, and serum short-chain fatty acids with fecal short-chain fatty acids. Biosci. Microbiota Food Health 39(1), 11–17. doi: 10.5281/zenodo.1439555).

Yang, Q., Liang, Q., Balakrishnan, B., Belobrajdic, D.P., Feng, Q.J., and Zhang, W. (2020). Role of Dietary Nutrients in the Modulation of Gut Microbiota: A Narrative Review. Nutrients 12(2). doi: 10.3390/nu12020381.

Zhang, X., Chen, W., Ning, Z., Mayne, J., Mack, D., Stintzi, A., et al. (2017). Deep Metaproteomics Approach for the Study of Human Microbiomes. Anal. Chem. 89(17), 9407–9415. doi: 10.1021/acs.analchem.7b02224.

Zhang, X., and Figeys, D. (2019). Perspective and Guidelines for Metaproteomics in Microbiome Studies. J. Proteome. Res. 18(6), 2370–2380. doi: 10.1021/acs.jproteome.9b00054.

Zhang, X., Li, L., Butcher, J., Stintzi, A., and Figeys, D. (2019). Advancing functional and translational microbiome research using meta-omics approaches. Microbiome 7(1), 154. doi: 10.1186/s40168-019-0767-6.

Zhang, X., Li, L., Mayne, J., Ning, Z., Stintzi, A., and Figeys, D. (2018). Assessing the impact of protein extraction methods for human gut metaproteomics. J. Proteomics 180, 120–127. doi: 10.1016/j.jprot.2017.07.001.

Zhang, X., Ning, Z., Mayne, J., Moore, J.I., Li, J., Butcher, J., et al. (2016). MetaPro-IQ: a universal metaproteomic approach to studying human and mouse gut microbiota. Microbiome 4(1), 31. doi: 10.1186/s40168-016-0176-z.

Zhong, H., Ren, H., Lu, Y., Fang, C., Hou, G., Yang, Z., et al. (2019). Distinct gut metagenomics and metaproteomics signatures in prediabetics and treatment-naive type 2 diabetics. EBioMedicine 47, 373–383. doi: 10.1016/j.ebiom.2019.08.048.

