## Supplemental Material for "Distinct human gut microbial taxonomic signatures uncovered with different sample processing and microbial cell disruption methods for metaproteomic analysis"

Supplementary Material

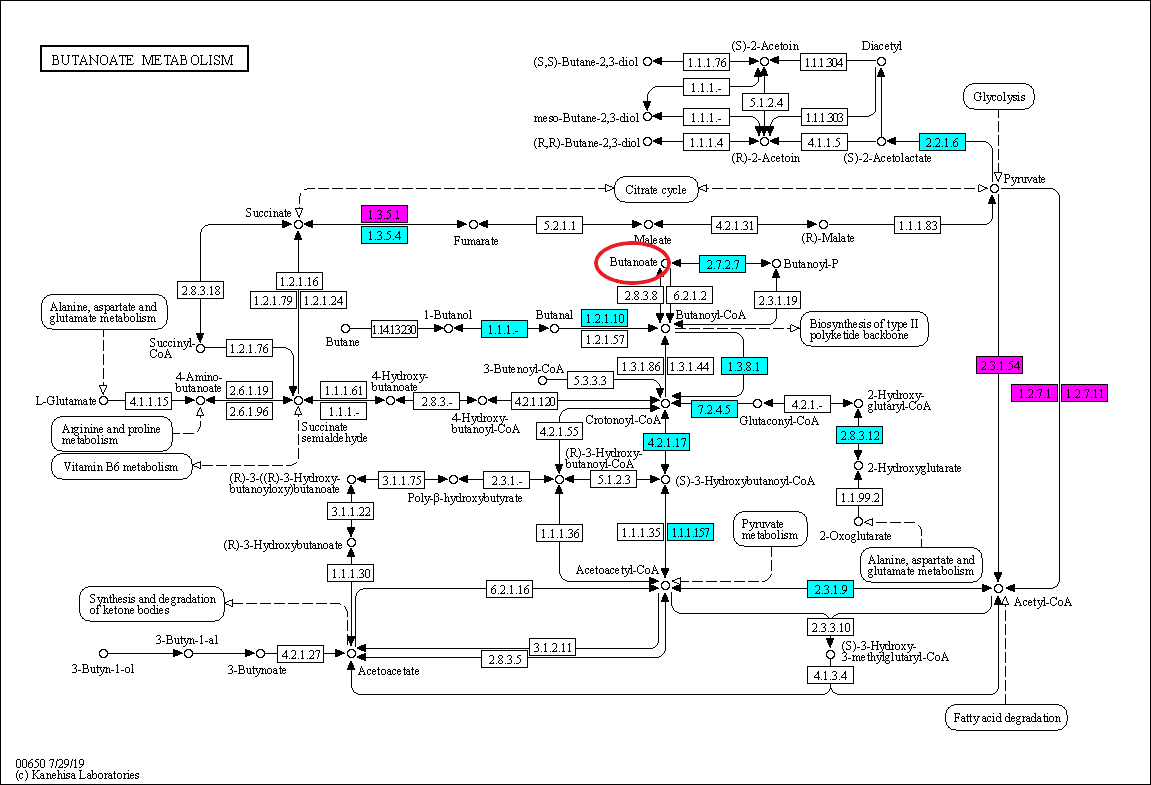

**Supplementary Figure 1.** Butyrate metabolism KEGG pathway downloaded from KEGG website (<http://www.kegg.jp>). Coloured KO numbers indicate proteins identified in this study. Firmicutes proteins are in coloured in blue and Bacteroidetes proteins in purple.
